# PKNs are required for cardiac hypertrophy induced by angiotensin II in mice

**DOI:** 10.64898/2026.10.26.761876

**Authors:** Susanna TE Cooper, Jacqueline JT Marshall, Peter J Parker, Angela Clerk

**Affiliations:** Institute of Developmental and Regenerative Medicine, Department of Physiology, Anatomy and Genetics, University of Oxford, UK; Protein Phosphorylation Laboratory, Francis Crick Institute, 1 Midland Road, London NW1 1AT, UK; Kings College London, Guy’s Campus, London SE1 1UL, UK; School of Biological Sciences, University of Reading, Whiteknights Campus, Reading, RG6 2AS, UK

**Keywords:** Cardiac hypertrophy, protein kinase N, angiotensin II, echocardiography, speckle-tracking

## Abstract

Of the three PKN isoforms, only homozygous Pkn2 knockout in mice is embryonic lethal; mice with Pkn2 haploinsufficiency or Pkn1/3 double knockout (Pkn1/3DKO) have no overt abnormalities. In adult mice, Pkn1/2 are implicated in development of cardiac disease, but studies of Pkn3 in the heart are lacking. Our hypothesis was that a role for Pkn1/3 in adult hearts would be revealed by pathophysiological stress. Here, we report expression profiles of PKNs in cardiac cells, identifying *PKN2* as the dominant isoform with expression of *PKN3* largely restricted to endothelial/immune cells. To test the hypothesis, Pkn1/3DKO, heterozygous Pkn2 knockout (Pkn2Het) and wild-type mice were treated with 0.8 mg/kg/d AngII, and cardiac function/dimensions assessed using speckle-tracking echocardiography. Baseline assessment identified minor differences in the left ventricle (LV) basal region in Pkn2Het (not Pkn1/3DKO) *vs* wild-type mice. AngII induced similar increases in LV end diastolic mass (LVEDM) in wild-type and Pkn1/3DKO mice at 3 d. LVEDM increased further in wild-type mice by 7d, but the initial response in Pkn1/3DKO mice was lost whilst stroke volume and cardiac output declined. Thus, hypertrophy was initiated in Pkn1/3DKO mouse hearts but not sustained. AngII increased cardiac fibrosis in wild-type mice together with mRNAs for cardiomyocyte hypertrophy and matrix proteins; this was suppressed in Pkn1/3DKO mice. Changes in expression of mRNA markers for cardiac non-myocytes suggested this was associated with vascular and immune cell dysfunction. In conclusion, Pkn2 does not substitute for Pkn1/3 in developing cardiac hypertrophy, and Pkn1/3 have an independent role in AngII-induced cardiac hypertrophy.

## Introduction

Heart failure affects millions of people worldwide with hypertension being one of the most prevalent risk factors [1]. The increase in blood pressure increases the workload of the heart. Initially, the heart adapts and cardiomyocytes (the contractile cells of the heart) increase in size and myofibrillar content (i.e. hypertrophy) to accommodate workload demand. However, prolonged haemodynamic overload leads to a complex remodelling programme with vascular remodelling, fibrosis and inflammation and such maladaptation can eventually lead to heart failure [2]. Understanding the molecular pathways governing both adaptive and maladaptive responses to hypertension remain fundamental to identifying new therapeutic strategies for heart failure.

The three members of the protein kinase N (PKN) family (PKN1, PKN2 and PKN3) were first identified as protein kinase C (PKC) -related kinases (PRKs) [3]. The mechanism of activation of PKC superfamily members is complex but, unlike the PKCs, PKNs are activated by Rho family small GTPases that are widely recognised for their role in regulating the actin cytoskeleton. Consistent with this, PKNs are involved in regulating the cytoskeleton and intracellular architecture, but also influence gene expression and glucose metabolism. Effects of PKNs on cell migration and cell cycle progression implicate them in various diseases. Only PKN2 is vital for life and homozygote Pkn2 knockout in mice is embryonic lethal [4]. These mice die at approximately embryonic day 10 (E10) with cardiovascular and neural tube defects. Heterozygote Pkn2 knockout (Pkn2Het) mice are viable and fertile with no obvious developmental defects. Cardiomyocyte-specific Pkn2 knockout causes cardiac developmental defects and surviving mice develop heart failure [5]. Pkn2 appears to play a very specific role in cardiomyocyte proliferation and sculpting of the heart at ∼E10.5, regulating the cytoskeleton, cell migration and formation of the ventricular wall [6]. In contrast to Pkn2, mice lacking either or both Pkn1 and Pkn3 (Pkn1/3 double knockout; Pkn1/3DKO) are viable and fertile with no obvious abnormalities [4].

PKNs are implicated in disease in the adult heart. In patients, PKN1 is downregulated in hibernating myocardium and is one of several genes deleted in a 19p13.12/19p13.3 congenital syndrome that includes cardiac abnormalities [7, 8]. By contrast, PKN2 is implicated in development of atrial fibrosis associated with atrial fibrillation [9]. Interestingly, *PKN2* mRNA is upregulated in dilated cardiomyopathy whilst *PKN1* and *PKN3* transcripts are downregulated [5]. In mice, cardiomyocyte-specific expression of constitutively-active Pkn1 protects cardiomyocytes from ischaemia-reperfusion injury [10], whilst Pkn1 knockout exacerbates injury in an *ex vivo* model of ischaemia-reperfusion [11]. *In vivo*, inducible cardiomyocyte-specific knockout of both Pkn1 and Pkn2 (but not each independently) in adult mouse hearts compromises cardiac hypertrophy resulting from pressure overload from transverse aortic constriction or in a hypertension model with angiotensin II (AngII) [12]. Although this suggests there may be redundancy between the two isoforms, our studies of Pkn2Het mice indicated that Pkn2 is specifically required for adaptation to AngII [5]. Pkn1/2 are also involved in the response of cardiac non-myocytes to injury, and fibroblast-specific deletion of Pkn1/2 attenuates cardiac fibrosis in mouse models of myocardial infarction and heart failure with preserved ejection fraction (HFpEF) by preventing fibroblast differentiation into myofibroblasts [13].

PKN3 has largely been studied in relation to cancer [14, 15], with little information about its role in the heart other than the observation that *PKN3* transcripts are downregulated in dilated cardiomyopathy [5]. However, its role in angiogenesis and maintenance of vascular integrity has important implications for cardiac pathology. Our hypothesis was that, even though global knockout of Pkn1 and Pkn3 in mice has no phenotypic effects [4], these kinases are likely to play an important role in adaptation to pathophysiological stress. For the heart, this would be detected in conditions such as increasing workload. Here, we report cardiac cell-type expression of the PKNs and, using Pkn1/3DKO mice, demonstrate that Pkn1/3 are required to sustain cardiac hypertrophy in a model of AngII-induced cardiac hypertrophy. Thus, Pkn2 does not substitute for Pkn1/3 in developing cardiac hypertrophy, and Pkn1 and/or Pkn3 play an important independent role in cardiac adaptation to stress.

## Results

### Expression of PKN isoforms in the heart

To compare the expression profiles of PKN isoforms in different cardiac cells, we interrogated the Human Protein Atlas (HPA) database for single cell transcriptomics. *PKN2* mRNA was the most highly expressed isoform in all cardiac cells (**Figure 1A**, **Supplementary Table S1**). *PKN1* was not detected in cardiac vascular smooth muscle cells (VSMCs) and had limited expression in adipocytes. This contrasts with other tissues in which there was significant expression in these cell types (**Supplementary Figure S1**).

**Figure 1.**
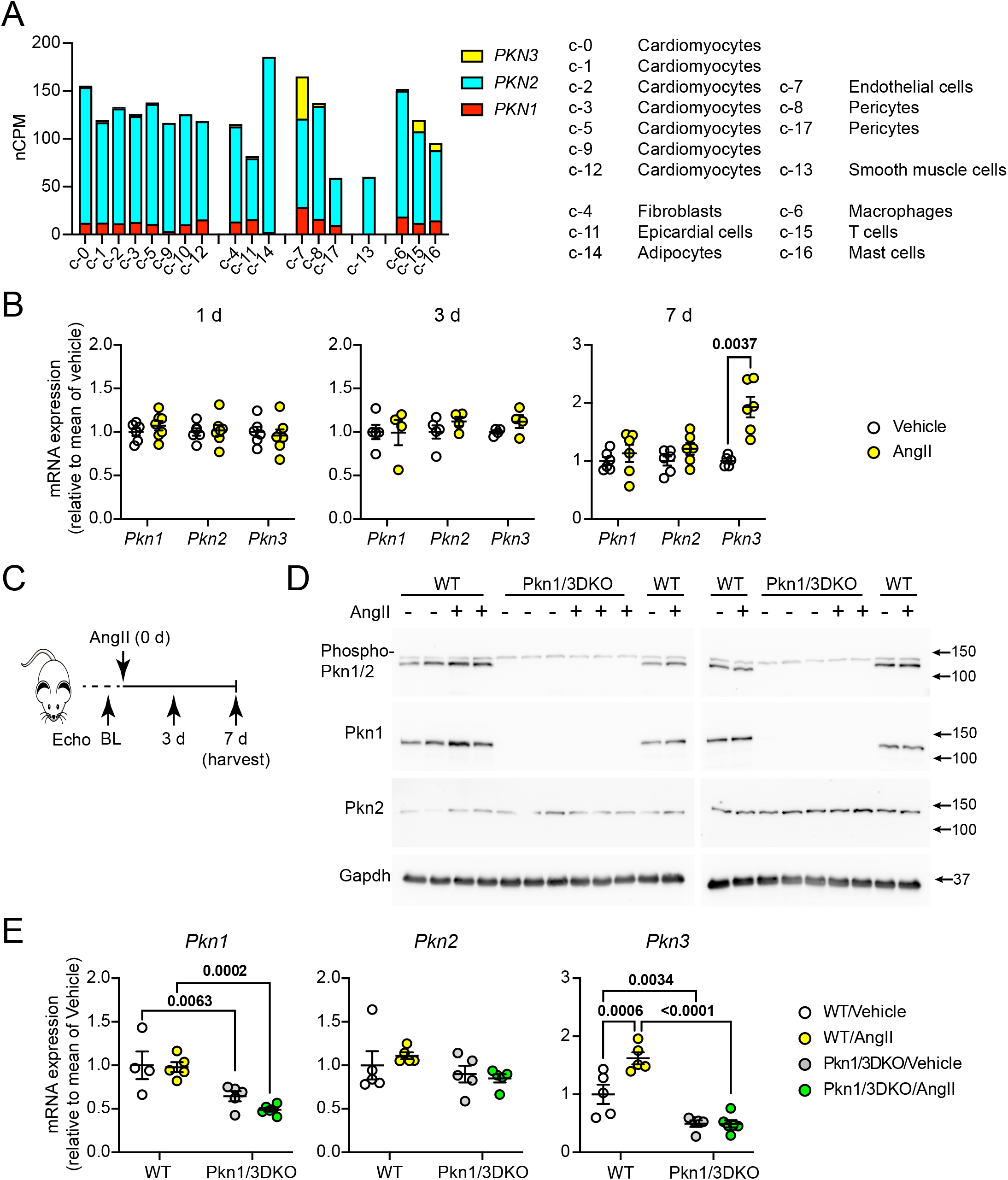
Expression of PKN isoforms in the heart. **A**, mRNA expression of *PKN1*, *PKN2* and *PKN3* in 17 cell types (c-0 to c-17) identified in human heart tissue. Data are from the Human Protein Atlas version 25.0. **B**, qPCR for *Pkn1*, *Pkn2* and *Pkn3* mRNA expression in hearts from male C57Bl/6J mice (10 wks) treated with vehicle or 0.8 mg/kg/d angiotensin II (AngII) for 1 d (left panel), 3 d (centre panel) or 7 d (right panel). Individual points are shown with means ± SEM. Statistical analysis used multiple unpaired t tests with Holm-Sidak’s multiple comparisons test. **C**, Schematic for *in vivo* experiments. Pkn1/3 double knockout (Pkn1/3DKO) or wild-type (WT) male mice (12 wks) were treated with vehicle or 0.8 mg/kg/d AngII for 7 d. Echocardiograms were collected prior to treatment (baseline, BL) and 3 or 7 d following the start of treatment. Hearts were harvested at 7 d. **D**, Expression of phosphorylated (phospho-) Pkn1/2 (upper images), total Pkn1 (images second from top), total Pkn2 (images second from bottom) and Gapdh (lower images) protein was measured by immunoblotting. Samples were from WT or Pkn1/3DKO mice treated with vehicle or AngII (7 d). **E**, qPCR for *Pkn1*, *Pkn2* and *Pkn3* mRNA expression in hearts of WT or Pkn1/3DKO mice treated with vehicle or AngII for 7 d. Individual points are shown with means ± SEM. Statistical analysis used 2-way ANOVA with Tukey’s post-test.

Expression of *PKN1* in other cardiac cells was 14.3 ± 6.9% (mean ± SD) that of *PKN2*, with highest relative expression in vascular endothelial cells (31.1%). Expression of PKN1 varied in different tissues, and the relative levels of PKN2 and PKN1 in cardiac cells in general was of a similar order as that of most other cells (**Supplementary Figure S2; Supplementary Spreadsheet SS1**). *PKN3* was predominantly expressed in vascular endothelial cells, with some expression in T cells and mast cells; other cardiac cells had little or no expression of *PKN3*. Using qPCR, we identified *Pkn3* as the only isoform upregulated in hearts from mice treated with 0.8 mg/kg/d AngII for 7 d (**Figure 1B**), suggesting that Pkn3 may be involved in disease progression.

### Global gene deletion of Pkn1 and Pkn3 in mice has a minimal effect on cardiac function and dimensions in mice

*PKN1* and *PKN3* mRNAs are downregulated in LV samples from dilated cardiomyopathy patients [5] suggesting that loss of both may be linked to cardiac disease. We used Pkn1/3DKO mice to investigate the joint roles of these isoforms in the heart. These mice were maintained as a homozygote line, so we compared cardiac function/dimensions with a related colony of wild-type and Pkn2Het mice (the closest match for genetic background and breeding/housing conditions [5]) both at baseline (this section) and following treatment with AngII (see below). Responses to AngII show marked sex dependence, with female mice often exhibiting attenuation or modification of AngII-induced cardiovascular phenotypes compared with males, potentially because of oestrogen-dependent mechanisms [16–21].

Furthermore, in the younger population, hypertension is more prevalent in men than in women and one of four priorities for sex-specific cardiovascular disease prevention in Europe is to improve detection and management of hypertension in men [22, 23]. To reduce biological variability and focus on the established male phenotype, only male mice were used in this study.

Following a baseline echocardiogram, mice were treated with 0.8 mg/kg/d AngII for 7 d (**Figure 1C**). Western blots confirmed deletion of Pkn1 protein with no effect on expression of Pkn2 (**Figure 1D**). Antibodies to detect endogenous Pkn3 in mouse samples could not be identified, so qPCR was used to compare mRNA expression levels. Transcripts of both *Pkn1* and *Pkn3*, but not *Pkn2* were reduced ∼50% in the hearts of Pkn1/3DKO mice compared with wild-type mice and, although *Pkn3* mRNA was upregulated in wild-type mouse hearts, there was no increase in *Pkn3* transcripts in AngII-treated Pkn1/3DKO mouse hearts (**Figure 1E**). Complete loss of *Pkn1/3* transcripts is not necessarily expected using qPCR methodology because transcription continues in a knockout model even if translation is aborted due to the presence of a STOP codon, and reduction in transcript expression results from nonsense-mediated decay. Importantly, there was comparable reduction in *Pkn1* and *Pkn3* transcripts matched by undetectable levels of Pkn1 protein.

Echocardiography with speckle-tracking (strain) analysis of long axis views of the left ventricle (LV) was used to determine if there were baseline cardiac abnormalities in Pkn1/3DKO mice. End diastolic volume (EDV) of Pkn1/3DKO mice was greater than wild-type mice, but there were no significant differences in end systolic volume (ESV) or LV end diastolic mass (LVEDM), nor were there any differences in EDV, ESV or LVEDM in Pkn2Het mice compared with wild-type mice (**Figure 2A-B**; **Supplementary Table S3**). Heart rates for Pkn1/3DKO or Pkn2Het mice were not statistically significantly different from wild-type mice, but there was a difference between Pkn1/3DKO and Pkn2Het mice that appeared to result from a small increase in Pkn2Het mice and a small decrease in Pkn1/3DKO mice (**Figure 2C**). The increase in EDV in Pkn1/3DKO mice was reflected in a small, non-significant increase in stroke volume, but (with a small decrease in heartrate), there was minimal effect on cardiac output. There were no differences in fractional shortening, ejection fraction or global longitudinal strain (GLS) between any of the mouse lines. Thus, there may be some effects of Pkn isoform knockout on cardiac function at baseline, though these appear minor. To explore this further, we assessed endocardial wall movement in different segments around the LV (segments 1 and 4 in the basal region, segments 2 and 5 in the mid-region and segments 3 and 6 at the apex), measuring peak radial and longitudinal displacement, velocity, strain and strain rate (see our recent characterisation of the model [24]) (**Figure 2D**). There were some significant differences in endocardial movement between Pkn2Het and wild-type or PKN2Het and Pkn1/3DKO hearts particularly in the basal regions of Pkn2Het mice, but we did not observe any significant differences between wild-type and Pkn1/3DKO mouse hearts (**Figure 2E-G**). We conclude that the hearts of Pkn1/3DKO mice were essentially normal.

**Figure 2.**
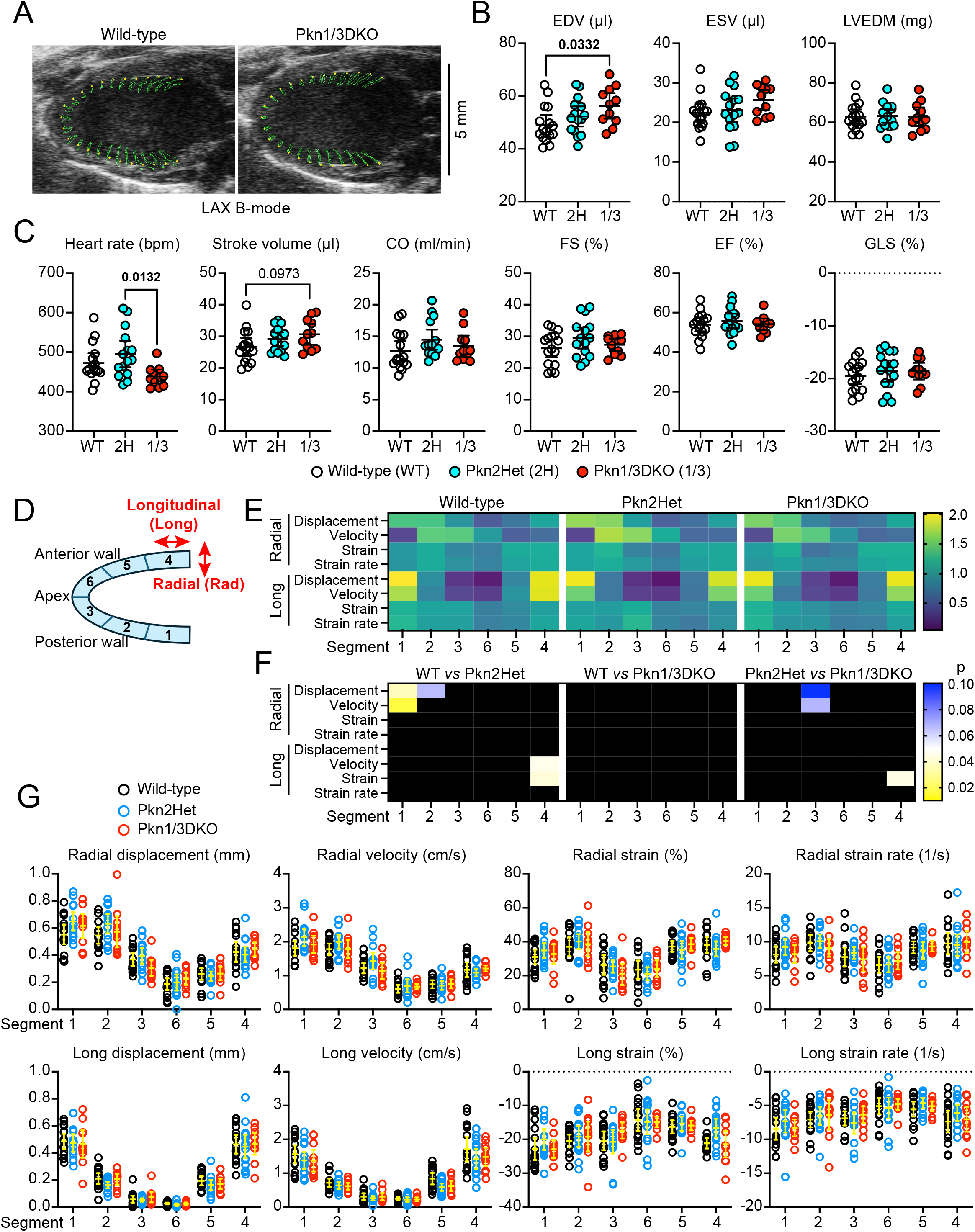
PKN1/3 gene deletion has a minimal effect on cardiac function and dimensions. Echocardiograms were collected from wild-type (WT), heterozygous Pkn2 knockout (Pkn2Het; 2H) and Pkn1/3 double knockout (Pkn1/3DKO; 1/3) male mice (12 wks). **A**, Representative B-mode long axis images showing the movement of the endocardial wall across two heartbeats traced in green from the yellow spots. **B-C**, Global speckle-tracking (strain) measurements from analysis of B-mode images from the long axis. **B**, Cardiac dimensions: end diastolic volume (EDV), end systolic volume (ESV), left ventricle end diastolic mass (LVEDM). **C,** Cardiac function: heart rate, stroke volume, cardiac output (CO), fractional shortening (FS), ejection fraction (EF), global longitudinal strain (GLS). **D**, Schematic diagram showing the positions of segments around the LV and orientation of longitudinal and radial movement (parallel and perpendicular to the endocardial wall, respectively). **E**, Heatmaps for each segment around the LV for wild-type (left panel), Pkn2Het (centre panel) or Pkn1/3DKO (right panel) mice showing mean peak values for radial and longitudinal (long) displacement, velocity, strain and strain rate (upper). **F**, Heatmaps for statistical analysis of segmental data showing p values for comparisons between WT and Pkn2Het mice (left panel), WT and Pkn1/3DKO mice (centre panel) and Pkn2Het and Pkn1/3DKO mice. **G**, Scatter plots for peak values for radial (upper graphs) and longitudinal (long; lower graphs) displacement, velocity, strain and strain rate. **B, C** and **G**, Individual points are shown with means ± 95% CI. The data meet conditions of normality testing; statistical analysis used one-way ANOVA with Tukey’s post-test. Significant differences (p<0.05) are in bold type.

### Loss of Pkn1/3 compromises AngII-induced cardiac hypertrophy

To determine if Pkn1/3 are likely to play a role in cardiac disease, we used a short-term (7 d) model of cardiac hypertrophy induced by 0.8 mg/kg/d AngII. This is a slow pressor dose that increases blood pressure over 7-14 d [25–27], but AngII also directly affects cardiac cells via AT1 receptors [28]. By 7 d, AngII significantly increased heart weight:body weight ratio in wild-type and Pkn2Het mice, consistent with cardiac hypertrophy, but this was not detected in Pkn1/3DKO mice (**Figure 3A**).

**Figure 3.**
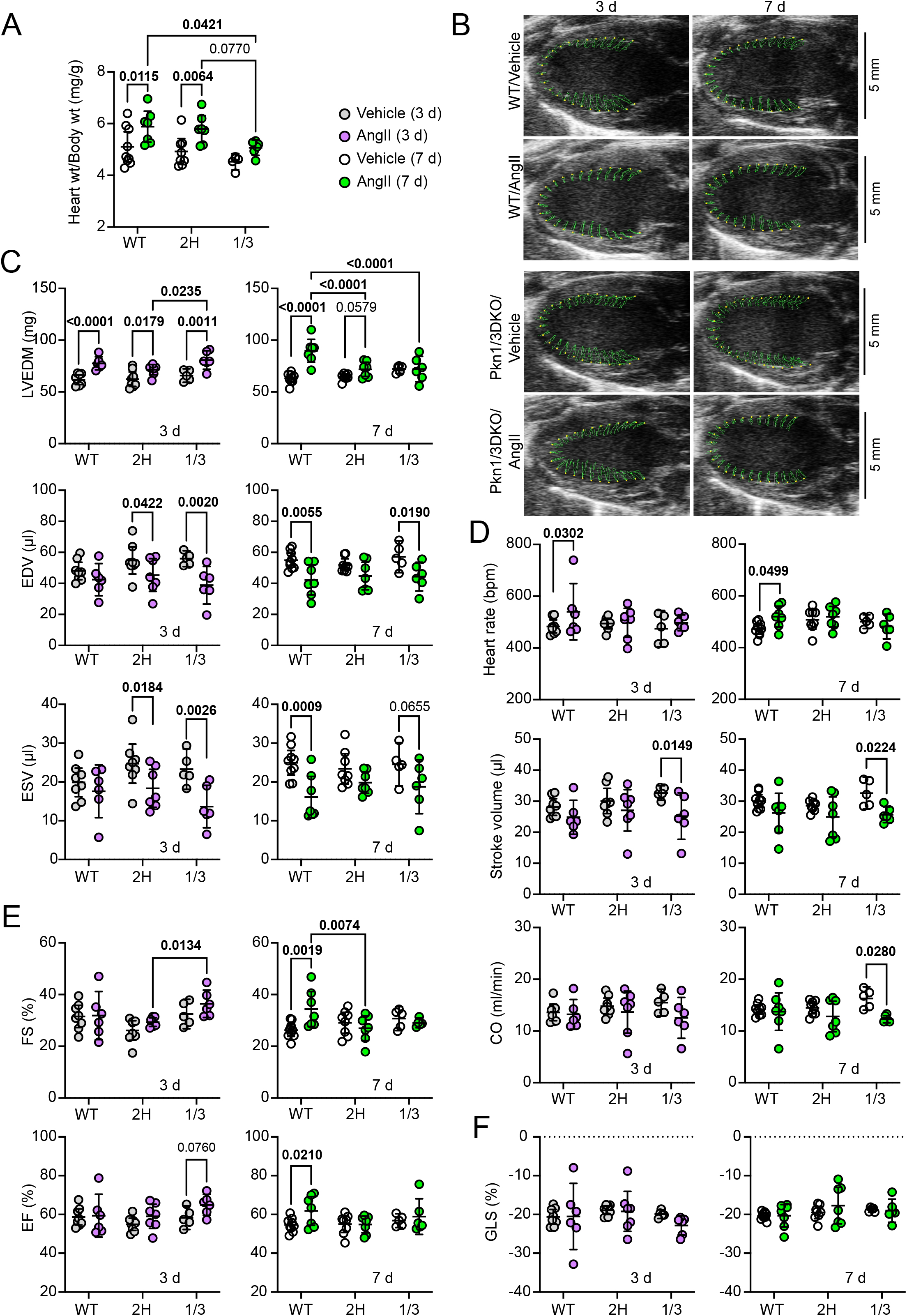
The hypertrophic response to angiotensin II is compromised in hearts from mice with heterozygous Pkn2 gene deletion or with Pkn1/3 double knockout compared with wild-type mice. Wild-type (WT), heterozygous Pkn2 knockout (2H) and Pkn1/3 double knockout (Pkn1/3DKO; 1/3) mice were treated with vehicle or 0.8 mg/kg/d angiotensin II (AngII) for 7 d. Long axis B-mode images were captured at 3 and 7 d. A, Heart weight (wt) to body wt ratio of hearts collected from mice treated with vehicle or AngII (7 d). B, Representative B-mode traces for WT or Pkn1/3DKO mice. Images at 3 and 7 d are from the same mouse. The movement of the endocardial wall across two heartbeats is traced in green from the yellow spots. C-F, Global strain measurements for WT, Pkn2Het and Pkn1/3DKO mice. C, Cardiac dimensions: end diastolic volume (EDV), end systolic volume (ESV), left ventricle end diastolic mass (LVEDM). D, Heart rate, stroke volume and cardiac output (CO). E, Fractional shortening (FS) and ejection fraction (EF). F, Global longitudinal strain (GLS). A, and C-F, Individual points are shown with means ± 95% CI. The data meet conditions of normality testing. Statistical analysis used two-way ANOVA with Tukey’s post-test with testing across 3 and 7 d data. Significant differences (p<0.05) are in bold type.

Cardiac dimensions were assessed using speckle-tracking of long axis B mode images taken at 3 and 7 d (**Figure 3B-F**; **Supplementary Table S4**). At 3 d, AngII increased LVEDM in wild-type, Pkn2Het and Pkn1/3DKO mice treated with AngII, but the response of Pkn1/3DKO mice was significantly greater than PKN2Het mice (**Figure 3C**, upper panels).

AngII induced a greater increase in LVEDM at 7 d in wild-type mice, but there was no further increase in Pkn2Het mice and the increase in LVEDM seen in Pkn1/3DKO mouse hearts at 3 d was lost (wild-type, 3 d *vs* 7 d: p=0.0038; Pkn2het 3 d *vs* 7 d: p=0.9288; Pkn1/3DKO, 3 d *vs* 7 d: 0.0408; two-way ANOVA with Tukey’s post-test). EDV and ESV were significantly decreased in hearts from Pkn2Het and Pkn1/3DKO mice treated with AngII for 3 d, whilst a significant effect was only detected at 7 d in wild-type mouse hearts (**Figure 3C**, centre and lower panels). Thus, Pkn2 haploinsufficiency and Pkn1/3 knockout each suppress AngII-induced LV hypertrophy, but in different ways: Pkn2 haploinsufficiency reduced the degree of hypertrophy which did not recover whereas hypertrophy was initiated in Pkn1/3DKO mice which then degenerated. For both, the earlier effect on EDV/ESV was likely to have an impact on cardiac function.

AngII induced a small increase in heart rate in wild-type mice, but cardiac output was not changed and, although there may have been some reduction in stroke volume, this was not significant (**Figure 3D**). Ejection fraction and fractional shortening were not significantly changed by AngII in wild-type mice with 3 d treatment, but both were increased at 7 d (**Figure 3E**), in line with the increase in LVEDM and decrease in EDV/ESV (**Figure 3C**).

Thus, wild-type mice adapted to the increased stress from AngII treatment. Despite the reduction in the AngII-induced increase in LVEDM and decreases in EDV/ESV, there were no significant effects of AngII on cardiac function Pkn2Het mice (**Figure 3D-E**). Interestingly, stroke volume and cardiac output were significantly inhibited in hearts from Pkn1/3DKO mice treated with AngII compared with vehicle, presumably because of the reduction in EDV/ESV in the absence of any significant changes in heart rate, fractional shortening or ejection fraction. There was no significant effect of AngII on GLS in any of the genotypes at either 3 or 7 d (**Figure 3F**). Overall, Pkn1/3 knockout had a greater effect on cardiac function than Pkn2 haploinsufficiency in mice treated with AngII.

Treatment of wild-type mice with 0.8 mg/kg/d AngII for 7 d promotes an increase in cardiomyocyte cross-sectional area (CSA), interstitial and perivascular fibrosis, increases in cardiac stress and hypertrophic mRNA markers (*Nppa*, *Myh7*, *Nppb*) alongside increases in expression of mRNAs for fibrosis-associated genes (e.g. *Col1a1*, *Col3a1*, *Fn1*, *Postn*) (**Figure 4**; and as shown in [5]). All of these changes induced by AngII were reduced in hearts from Pkn2Het mice (previously reported [5]). CSA of cardiomyocytes from Pkn1/3DKO mice appeared greater than that of wild-type mice, but AngII did not increase this further (**Figure 4A-B**). The increase in *Nppa* induced by AngII was similar in Pkn1/3DKO and wild-type mouse hearts, but the increase in expression of the hypertrophic markers *Myh7* and *Nppb* was significantly reduced (**Figure 4C**). AngII increased interstitial and perivascular fibrosis in wild-type mouse hearts, both of which were reduced in Pkn1/3DKO hearts (**Figure 4D-F**). Consistent with the reduction in fibrosis, expression of mRNAs for fibrosis-associated genes (*Col1a1*, *Col3a1*, *Fn1*, *Postn*) was reduced in hearts from Pkn1/3DKO mice relative to wild-type mice (**Figure 4G**). The data indicate that Pkn1/3 are required for cardiomyocyte hypertrophy and cardiac fibrosis.

**Figure 4.**
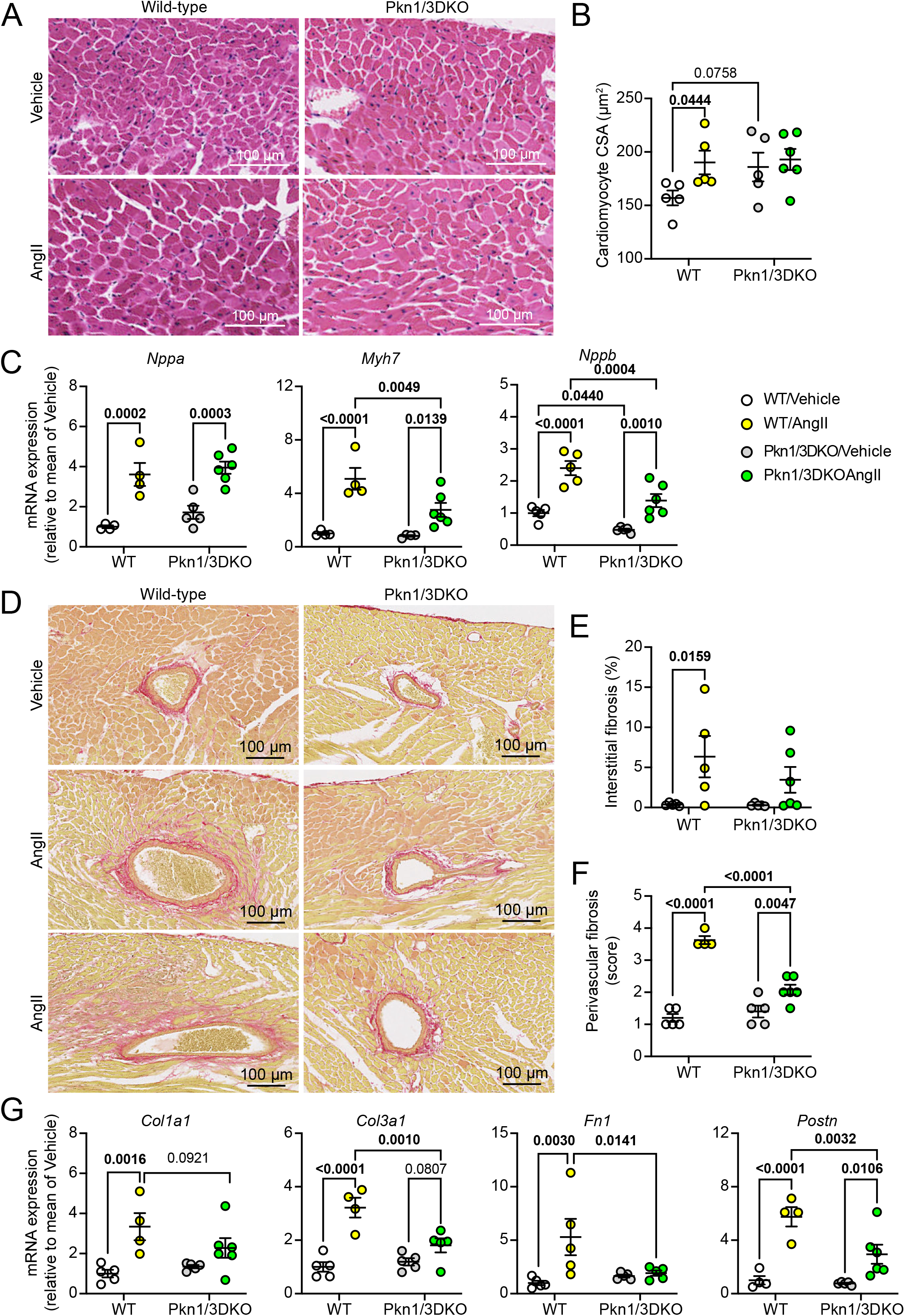
Pkn1/3DKO mice have reduced cardiomyocyte hypertrophy and cardiac fibrosis in response to angiotensin II compared with wild-type mice. Wild-type (WT) and PKN1/3 double knockout (Pkn1/3DKO) mice were treated with vehicle or 0.8 mg/kg/d angiotensin II (AngII) for 7 d. Hearts were harvested and fixed for histological analysis or snap frozen for RNA extraction and qPCR. A, Representative images of sections from wild-type (WT) and Pkn1/3DKO mice stained with haemotoxylin and eosin (H&E). B, Cardiomyocyte cross-sectional area (CSA) from H&E-stained sections. C, qPCR for expression of cardiomyocyte stress and hypertrophy markers. D, Representative images of sections from WT and Pkn1/3DKO mice stained with picrosirius red. Representative average images are shown in the upper and centre panels with the most severe examples in the lower panels. **E-F**, Measurement of interstitial (**E**) and perivascular (**F**) fibrosis. For interstitial fibrosis the area of fibrosis is presented as a percentage of the whole area of the myocardium. For perivascular fibrosis, the degree of fibrosis was scored: 0.5, small amount of fibrosis around a single vessel; 1, small amount of fibrosis around 2 or more vessels; 1.5, significant fibrosis around a single vessel; 2, significant fibrosis around 2 or more vessels; 2.5, extensive fibrosis around 1 vessel; 3, extensive fibrosis around 2 or more vessels; 3.5 extensive fibrosis around a single vessel penetrating into the myocardium with extensive fibrosis around other vessels; 4, extensive fibrosis around 2 or more vessels penetrating into the myocardium. **G**, qPCR for expression of selected markers of fibrosis. **B-C** and **E-G**, Individual points are shown with means ± SEM. Statistical analysis used 2-way ANOVA with Tukey’s post-test; significant differences (p<0.05) are in bold type.

### Loss of Pkn1/3 is associated with changes in mRNA cell markers for cardiac non-myocytes in AngII-induced cardiac hypertrophy

For further insight into the changes associated with the compromised hypertrophic response to AngII in Pkn1/3DKO mouse hearts, we used qPCR for mRNA cell markers to assess potential changes in the cellular composition of cardiac non-myocytes. In humans, *PKN3* is predominantly in endothelial cells in the heart which also have relatively high expression of *PKN1*. Thus, by extrapolation to mouse hearts, Pkn1/3 may be particularly important in their response to AngII. Consistent with this, the increase in expression of endothelial cell markers *Pecam1* (*Cd31*, [29]) and *Sele* (E-selectin, [30]) induced by AngII in wild-type mouse hearts was inhibited in Pkn1/3DKO mice (**Figure 5A**, left two panels). Pericytes are required to maintain blood vessel integrity [31], but do not express *PKN3* and have relatively low expression of *PKN1* in human cells (**Supplementary Table S1**). AngII increased expression of the pericyte marker *Cspg4* (*Ng2*) in wild type mouse hearts (**Figure 5A**, centre panel), consistent with an increased requirement for pericytes in association with endothelial cells. This was not inhibited in Pkn1/3DKO mouse hearts, but baseline expression was already significantly higher in hearts from vehicle-treated Pkn1/3DKO mouse hearts compared with wild-type hearts. We previously reported that *Tagln* mRNA (*Sm-22a,* expressed in VSMCs and early-stage cardiomyocytes [32]) is upregulated in wild-type mouse hearts following AngII treatment [5]. Although VSMCs only appeared to express *PKN2* in human hearts (**Figure 1A**), AngII-induced expression of *Tagln* was inhibited in Pkn1/3DKO mouse hearts (**Figure 5A**, second to right panel). Together with the endothelial cell markers, this suggests there is likely to be vascular dysfunction. *Ddr2* is considered a marker of fibroblasts [33] and, although there was reduced fibrosis in hearts from Pkn1/3DKO mice treated with AngII compared with those from wild-type mice (**Figure 4D-G**), there were no significant changes in Ddr2 expression (**Figure 5A**, right panel) suggesting that the number of these fibroblasts was not a factor in the fibrotic response.

**Figure 5.**
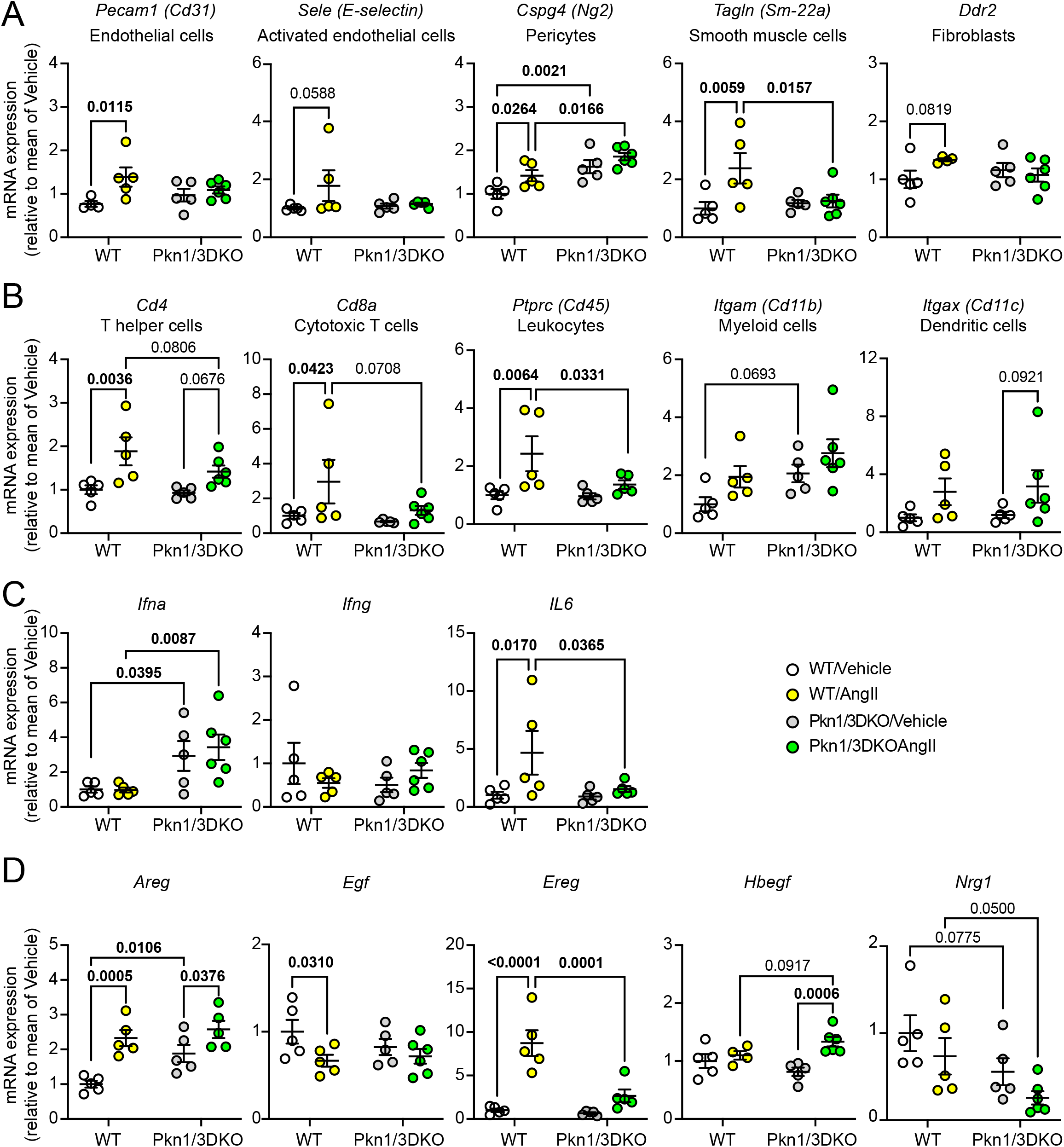
mRNA expression of cell-specific and immune markers, and epidermal growth factor (EGF) receptor family ligands in hearts from Pkn1/3DKO mice relative to wild-type mice. Wild-type (WT) and PKN1/3 double knockout (Pkn1/3DKO) mice were treated with vehicle or 0.8 mg/kg/d angiotensin II (AngII) for 7 d. Hearts were harvested and snap frozen for RNA extraction and qPCR for selected genes. A, mRNA expression of gene markers for cardiac non-myocytes. B, mRNA expression of gene markers for cardiac immune cells. C, Expression of *Ifna*, *Ifng* and *IL6* mRNAs. D, Expression of EGF receptor ligands. Individual points are shown with means ± SEM. Statistical analysis used 2-way ANOVA with Fisher’s LSD tests; significant differences (p<0.05) are in bold type.

Cardiac hypertrophy and heart failure in pressure-overload conditions (including AngII-dependent mouse models) is associated with inflammation and infiltration of immune cells [34, 35]. Since *PKN3* was expressed in some cardiac immune cells (**Figure 1A**), we considered if there were effects on the immune response to AngII. Consistent with the relatively high levels of *PKN1* and *PKN3* in T cells in humans, *Cd4* and *Cd8a* mRNAs (markers of helper and cytotoxic T cells, respectively) were significantly increased by AngII in hearts from wild-type mice, but not in hearts from Pkn1/3DKO mice (**Figure 5B**, left two panels). A similar profile was seen for *Ptprc* (*Cd45*), a general leukocyte marker [36] (**Figure 5B**, centre panel). However, other immune cells were not affected by Pkn1/3DKO and expression of *Itgam (Cd11b*, a marker of myeloid cells including monocyte/macrophages that infiltrate the heart with AngII infusion [37]) and *Itgax* (*Cd11c*; a marker of dendritic and other immune cells [38]) were increased to some degree by AngII with similar profiles in wild-type and Pkn1/3DKO mouse hearts (**Figure 5B**, right two panels). Further indication of dysregulation of the immune response was the significant increase in expression of *Ifna* (though not *Ifng*) mRNA in Pkn1/3DKO mouse hearts, independent of AngII-treatment, whilst the increase in expression of IL6 induced by AngII in wild-type mouse hearts was inhibited in Pkn1/3DKO mouse hearts (**Figure 5C**). The data indicate that Pkn1/3 contribute to selected aspects of AngII-induced inflammation in mouse hearts.

Changes in expression of mRNA markers suggest that loss of Pkn1/3 affects the cellular composition of the heart and the immune response; these could reflect loss of Pkn1/3 within cardiac cells and/or result from effects on intercellular mediators and autocrine/paracine signalling. To determine if the latter may be a factor, we assessed mRNA expression of ligands for the EGF receptor family (Egfr, Erbb3, Erbb4) that are subject to regulation in hearts of mice treated with AngII [39] (**Figure 5D**). Some ligands recognise only Egfrs (e.g. Areg, Egf) whilst others bind to Egfrs or Erbb3 (e.g. Ereg, Hbegf) and neuregulins (e.g. NRG1) bind to Erbb3 and Erbb4 . AngII increased expression of *Areg* mRNA in wild-type and Pkn1/3DKO mouse hearts but, because *Areg* mRNA was already upregulated in Pkn1/3DKO mouse hearts, the further increase with AngII was reduced.

Expression of *Egf* mRNA was decreased in hearts from wild-type mice treated with AngII, an effect that was not seen in Pkn1/3DKO mouse hearts, probably because expression levels already appeared to be reduced. *Ereg* mRNA was substantially upregulated by AngII in wild-type mouse hearts and this was suppressed in Pkn1/3DKO mouse hearts, whilst *Hbegf* mRNA expression was increased by AngII in Pkn1/3DKO but not wild-type, mouse hearts.

*Nrg1* mRNA was downregulated in hearts from Pkn1/3DKO mice treated with AngII. Thus, Pkn1/3 knockout causes dysregulation of the EGF receptor family network, an example of how intercellular signalling in the heart may be disrupted through loss of these kinases.

## Discussion

The PKNs were identified as PKC-related kinases over 30 years ago [40–42], but much remains to be discovered about their mechanisms of regulation, their functional roles and how they contribute to disease. All three isoforms are expressed in the adult heart (**Figure 1**), but only Pkn2 knockout is embryonic lethal in mice, with no obvious phenotype in mice with global knockout of Pkn1, Pkn3 or both [4]. This could indicate functional redundancy, with Pkn2 being sufficient to compensate for loss of Pkn1/3 (or complete redundancy of Pkn1/3), but Pkn1/3 could still be needed for adaptation to pathophysiological stresses, the hypothesis tested here. Using echocardiography, we confirmed that hearts from adult male Pkn1/3DKO mice were essentially normal, although there were minor effects of Pkn2 haploinsufficiency (**Figure 2**). In support of our hypothesis, the hearts of Pkn1/3DKO mice failed to respond normally to the imposition of an extracellular stress, AngII (**Figure 3-4**).

Interestingly, the response of Pkn1/3DKO mouse hearts to AngII was dissimilar to that of Pkn2Het mouse hearts and, for example, Pkn2 haploinsufficiency inhibited the increase in LVEDM induced by AngII in wild-type mice at 3 and 7 d, whereas hearts from Pkn1/3DKO mice showed a comparable increase in LVEDM as in wild-type mice at 3 d which regressed by 7 d. This suggests that cardiac adaptation was initiated in the absence of Pkn1/3, but the process subsequently failed. The mechanisms require further investigation, but the effects on mRNA markers for cardiac non-myocytes suggest a link to vascular dysfunction and/or immune regulation, potentially with dysfunction of autocrine/paracrine signalling (**Figure 5**). Irrespective of mechanism, the data suggest that there is not functional redundancy between Pkn2 and the other isoforms.

### PKN expression profiling in cardiac cells

We and others have studied PKN expression in the heart and in cardiomyocytes, but this is generally at the whole tissue level or in cardiomyocytes and is not as informative as single cell transcriptomics that can distinguish between multiple populations of cardiomyocytes and identify cardiac non-myocytes. We therefore used the HPA database for comprehensive cardiac cell mapping of PKN mRNA expression. As in most other tissues, *PKN3* mRNA was almost exclusively expressed in cardiac endothelial cells, with some expression in T cells and mast cells, whilst *PKN2* was the dominant isoform expressed in all cardiac cells (including cardiac endothelial cells) (**Figure 1A**, **Supplementary Table S1**, **Supplementary Spreadsheet SS1**). *PKN1* expression was more varied. Cardiac VSMCs were the only VSMCs with apparently no *PKN1* mRNA, raising the question of why those in the heart appear to be anomalous. Expression of *PKN2* was also low in cardiac VSMCs compared with those from other tissues, so it could be a technical issue or, perhaps, cardiac VSMCs (presumably in arterioles in the heart) are more specialised. The situation is similar for cardiac adipocytes with extremely low *PKN1* mRNA compared with those in other tissues.

PKN1 transcripts were expressed in all cardiomyocyte populations but at a substantially lower level than those for *PKN2* with an average ratio of *PKN2:PKN1* of 12.5:1, but this was variable (range 6.5-32.3:1).

With any transcriptomics data, the question remains of relative protein levels. This is difficult to address on a comparative level. We and others have used immunoblotting to demonstrate expression of Pkn1 and Pkn2 in rodent hearts or cardiomyocytes but this technique is generally semi-quantitative and antibodies are not available that recognise endogenous levels of Pkn3. Proteomics analysis is challenging for protein kinase studies in cardiomyocytes (and therefore the heart) because expression levels of kinases are low compared with other proteins required for, for example, the contractile apparatus or mitochondria. It is possible to increase the sensitivity for kinase detection by first enriching for protein kinases prior to identification by mass spectrometry. Data for cardiomyocytes indicate 9.6-fold greater expression of Pkn2 than Pkn1 in neonatal rat cardiomyocytes and 2.4-fold in adult rat cardiomyocytes, with no identification of Pkn3 [43]. Allowing for post-transcriptional regulation of protein expression, this is largely consistent with the HPA transcriptomics data. We are not aware of proteomics/kinomics studies for other cardiac cells.

### Cardiac adaptation to AngII in male mice and the Pkns

We, like others, used a model of cardiac hypertrophy in mice induced by AngII. However, different concentrations of AngII can have different effects on the heart with, for example, lower concentrations promoting diastolic dysfunction (effects more likely to result from direct action of AngII on AT1 receptors on cardiac cells) whilst higher concentrations have a greater effect on blood pressure leading to systolic dysfunction with reduced ejection fraction (i.e. effects due to increased workload on the heart) [44]. The dose used here is described as a slow-pressor dose [25–27] that increases blood pressure gradually over 7-14 d, a model with features of adaptive hypertrophy and sustained ejection fraction at early stages (up to 7 d) but which becomes maladaptive over 28 d with a significant decline in ejection fraction and GLS [45]. This study focused only on the early adaptive phase over 7 d, during which time direct effects on cardiac cells may be expected to predominate. Serial echocardiography is essential for understanding how mouse hearts respond to AngII and we have recently conducted a detailed analysis of this model using echocardiography with speckle-tracking and segmental strain, highlighting changes between 3 and 7 d and differences in response of different lines of mice in this early phase [24]. Here, we compared the response of Pkn1/3DKO mice to both wild-type and Pkn2Het mice on the same genetic background (C57Bl/6J) and bred in the same institution (i.e. the best comparator available). The responses of the wild-type mice were similar to those of C57Bl/6J mice from a commercial supplier, but not identical. For example, although both lines had a greater increase in LVEDM at 7 d compared with 3 d, wild-type mice in [24] had decreased EDV at 3 and 7 d (not just 7 d as in this study) with reduced cardiac output (cardiac output was maintained in this study).

AngII-induced cardiac hypertrophy (was suppressed in both Pkn2Het and Pkn1/3DKO mice, but the responses were different. Interestingly, there was significant suppression of overall cardiac hypertrophy (measured by heart weight: body weight ratio) at 7 d in Pkn1/3DKO mice but not Pkn2Het mice (**Figure 3A**). As mentioned above, the AngII-induced increase in LVEDM was inhibited in Pkn2Het mice from 3 d but, whilst adaptation over 3 d appeared normal in Pkn1/3DKO mice, this was lost by 7 d (**Figure 3C**), suggesting that adaptation was initiated but then failed. Assuming most of the initial adaptation results from direct effects of AngII on cardiac cells, this may reflect the ubiquitous expression of Pkn2 in all cells such that global Pkn2 haploinsufficiency compromises the overall response, compared with a more restricted role of Pkn1 and/or Pkn3 and a more limited effect of global knockout.

PKN3 expression predominates in endothelial cells, and it is implicated in angiogenesis in various models including tumour development [14]. Given its expression in cardiac endothelial cells (**Figure 1A**), it is likely that Pkn3 is required to support angiogenesis in the heart and Pkn3 knockout has the potential to compromise this response. Reduced angiogenesis is predicted to restrict cardiomyocyte hypertrophy as the pathology develops over time. Thus, the increase in Pkn3 mRNA expression in wild-type hearts from mice treated with AngII potentially reflects an increase in endothelial cells that cannot be accommodated in the Pkn1/3DKO mouse hearts (**Figure 1, B and E**). Furthermore, mRNA expression data support the concept that angiogenesis is compromised in Pkn1/3DKO mouse hearts, since endothelial cell markers induced by AngII in wild-type mouse hearts were not increased in hearts from Pkn1/3DKO mice (**Figure 5A**). Experiments with conditional deletion of Pkn2 *vs* Pkn3 in endothelial cells would be useful to understand their roles in these cells. *Tagln* mRNA, a marker of VSMCs in the adult heart was increased by AngII in wild-type mouse hearts (**Figure 5A**). The absence of a response in Pkn1/3DKO mouse hearts may be an indirect effect of the loss of the endothelial cell response and/or changes in the intercellular signalling, but this further suggests vascular dysfunction that is likely to compromise the development of cardiac hypertrophy.

The most simplistic view of cardiac adaptation to AngII treatment in the early phase is that it results from a direct effect of AngII on cardiac cells, given that all cardiac cells express receptors for AngII and many studies report that AngII can promote cardiomyocyte hypertrophy *in vitro* and *in vivo* [28]. However, exposure of cardiac cells to AngII immediately changes their responsiveness to the environment resulting from changes in gene expression. Our studies have focused mostly on other hypertrophic agents (e.g. endothelin-1, α_1_-adrenergic agonists such as phenylephrine), but such stimuli stimulate changes in immediate early genes within 15-30 min and changes in protein expression over the subsequent 1-2 h [46, 47]. These include transcriptional regulators and intercellular mediators such as the growth factors that activate the EGF receptor family as monitored here. These and other intercellular mediators are likely to operate in ever-changing autocrine/paracrine networks that potentiate cardiac hypertrophy induced by AngII, rather than AngII acting in isolation. Indeed, inhibition of the EGF receptor family reduces cardiomyocyte hypertrophy and enhances cardiac fibrosis in this AngII model [39].

Consistent with the data in Fuller et al. [39], AngII significantly increased *Ereg* mRNA expression with no significant change in *Hbegf* and both had a suggestion that there was some decrease in *Nrg1* in wild-type mouse hearts. However, although expression of *Areg* and *Egf* were upregulated and downregulated, respectively, in this study (**Figure 5D**), their expression was unchanged in Fuller et al. [39], possibly a consequence of differences between the mouse lines. Nevertheless, modulation of the changes in expression of these EGF receptor family ligands in Pkn1/3DKO mouse hearts indicates that autocrine/paracrine signalling is likely to be substantially altered compared with wild-type mouse hearts.

Consequently, although the changes in mRNA expression for cell markers are likely to reflect cellular content, these changes may not be directly due to loss of Pkn1/3 in those particular cells but could instead result from defective intercellular signalling.

### Cardiac adaptation to AngII in male and female mice

This study (like many others) focused exclusively on young male mice in which (since hypertension is more prevalent in young men [22, 23]) the AngII model has particular clinical relevance. Young female rodents are resistant to AngII-induced hypertension and cardiac hypertrophy with evidence to indicate that this is oestrogen dependent [16–21]. For example, ovariectomised young female mice become sensitised to AngII-induced hypertension, a response that is suppressed by oestrogen administration. Another factor is aging and older female mice also become more sensitive to AngII. The mechanisms are under investigation but may relate to depressor activity of AT2 receptors that, in females, attenuate the pro-hypertensive effect of AT1 receptors [18], and/or sex differences in NO production [48]. Differences between young male and female mice are apparent in other models of cardiac hypertrophy including a preclinical model of heart failure with preserved ejection fraction (HFpEF) and cardiac hypertrophy induced by the α_1_-adrenergic agonist, phenylephrine [49, 50]. The data strongly support the argument for preclinical studies in female animals in addition to males, but an alternative argument to be made is that preclinical studies in either sex should be linked more closely to human cardiovascular epidemiology and, for women, this extends to assessment of hormonal lifetime changes.

### Global gene deletion of PKNs in mice *vs* cell-specific knockout studies

In this and our related study of Pkn2 in the heart [5] we used genetically-altered mice with global gene deletion, whilst other researchers have studied Pkn1 and Pkn2 (though not Pkn3) using models for tamoxifen-inducible gene knockout in cardiomyocytes and cardiac fibroblasts [12, 13]. The studies with cardiomyocyte-specific knockout have proposed that Pkn1 and Pkn2 may be functionally redundant in these cells since knockout of both isoforms (Pkn1/2DKO) is necessary to prevent cardiac hypertrophy induced by transverse aortic constriction or AngII. Our studies suggest otherwise given that we observe subtly distinct phenotypes in Pkn2Het *vs* Pkn1/3DKO mice treated with AngII. One consideration is that effects of Pkn1 and/or Pkn2 knockout in an inducible cardiomyocyte-specific knockout model (even assuming full inducibility) are likely to have a more limited effect on cardiac hypertrophy *in vivo* than global knockout. This is because both isoforms are expressed in all cardiac cells and cardiomyocytes represent only ∼30% of the total cell number in the heart, the rest of the cells being predominantly endothelial cells and fibroblasts [51, 52].

Furthermore, loss of Pkn1 and Pkn2 in the Pkn1/2DKO model may reflect an additive or synergistic consequence of different functional effects rather than, necessarily, substitution of one for the other. An alternative consideration is that the kinases are functionally redundant, and the apparent non-redundancy in our studies is because we used a global knockout system but PKNs are not uniformly expressed in all cells (e.g. expression of Pkn3 in endothelial cells). Clearly, further studies are required.

Pkn1/2 seem to be particularly implicated in cardiac fibrosis. Thus, fibrosis is reduced in both cardiomyocyte- and fibroblast-specific Pkn1/2DKO mice in models of pressure overload (cardiomyocyte Pkn1/2DKO mice), myocardial infarction or HFpEF (fibroblast Pkn1/2DKO mice) [12, 13]. In fibroblasts, there is reduced differentiation into myofibroblasts and expression of extracellular matrix proteins in Pkn1/DKO mouse hearts. Our studies with Pkn2Het and Pkn1/3DKO mice are consistent with these studies since AngII-induced expression of mRNAs encoding extracellular matrix proteins and increased fibrosis were inhibited in both genotypes compared with wild-type mice, along with histological evidence of fibrosis [5] (**Figure 4F-G**).

### The immune system in AngII-induced cardiac hypertrophy

The importance of the immune system in cardiac pathology associated with myocardial infarction is well-recognised but it also plays a significant role in other cardiac pathologies including hypertension [34, 53–55]. Expression of *PKN3* mRNA in cardiac T cells (**Figure 1A**, **Supplementary Spreadsheet SS1**) raised the question of the effect of Pkn1/3DKO on AngII-induced changes in immune cell and cytokine production. Although limited in nature and restricted to mRNA expression (**Figure 5C-D**), the data in this study point to dysregulation of some, but not all aspects of the immune response in Pkn1/3DKO mouse hearts both at baseline (e.g. increased expression of *Ifna*) and in response to AngII (e.g. reduced expression of T cell markers). Because of the limitations associated with using global knockout mouse models, it is not possible to determine if the effects reflect a response of the cardiac immune system itself or may be a result of infiltrating immune cells. More extensive studies are needed to clarify the role of PKNs, particularly Pkn3, in the immune system in the heart and the wider context of the immune system in general.

### Conclusions

This study established the variation in PKN isoform expression between different cardiac cells and demonstrated that, even though global knockout of Pkn1 and Pkn3 in mice has no detectable effects on cardiac function or dimensions in the absence of a stressor, Pkn1/3 are required for cardiac hypertrophy induced by AngII in mice. Thus, loss of Pkn1/3 inhibited progression of AngII-induced cardiac hypertrophy, reduced cardiomyocyte hypertrophy and fibrosis, and caused cardiac dysfunction even in the short term. The data support the concept of distinct roles for Pkn1 and Pkn2 in the overall hypertrophic response, because Pkn2 does not substitute for Pkn1/3 in developing cardiac hypertrophy. Instead, Pkn1 and/or Pkn3 appear to operate independently of Pkn2 in cardiac adaptation to AngII.

## Methods

### mRNA expression of PKN isoforms in human cardiac cells

Single cell mRNA data for heart tissue were from the Human Protein Atlas (HPA) version 25.0 (release date: 11 November 2025) [56]. Cluster data from GSE183852 [57] were downloaded (23 April 2026) from https://www.proteinatlas.org/humanproteome/single+cell/single+cell+type/data#cluster_data and the data mined for expression of *PKN1*, *PKN2* and *PKN3*. Data are presented as normalised expression (nCPM).

### Ethics statement for *in vivo* studies

Mouse breeding was conducted at the Biological Research Facility at the Francis Crick Institute. Adult male mice (average age 11 wks) were transferred to the BioResource Facility at St. George’s University of London for experiments. Each site is UK registered with a Home Office certificate of designation. Procedures were performed in accordance with UK regulations and the European Parliament Directive 2010/63/EU for animal experiments.

Work was undertaken in accordance with local institutional animal care committee procedures and the U.K. Animals (Scientific Procedures) Act 1986. Experiments were conducted under Project Licence P8BAB0744.

### *In vivo* mouse experiments

Mice were maintained on a C57Bl/6J background and were sourced as described in [4]. PKN1/3DKO mice are viable and fertile with no obvious phenotype and breeding was conducted as a homozygote line. Wild-type mice were sourced from the Francis Crick Institute, the most closely related mice being those derived from breeding of the heterozygote Pkn2 line. Experiments with wild-type, heterozygote Pkn2 mice and Pkn1/3DKO mice were conducted between October 2020 and June 2021 by the same investigators. Experiments comparing Pkn2Het mice with wild-type littermates were previously reported in [5] and echocardiograms were re-analysed for this study. An additional study of wild-type mice treated with vehicle (n=4) or AngII (n=2) was conducted subsequently.

Mice were allowed to acclimatise for 7 d following transfer and randomly allocated to treatment groups. No mice were excluded after randomisation. Individuals conducting the studies were not blinded to experimental conditions for welfare monitoring purposes.

Animals were checked daily and mice undergoing procedures were monitored using a score sheet. Weights were taken before, during and at the end of the procedures (**Supplementary Table S5**). All mice are routinely culled if they reach a predefined endpoint agreed with the Named Veterinary Surgeon but no mice reached this point in the experiments included in this study. No other exclusion criteria were set *a priori* for the experimental process. Following minipump surgery or echocardiography, mice were recovered singly and returned to a clean cage once fully recovered. At the end of the experiment, mice were culled by cervical dislocation with severance of the femoral artery to ensure cessation of life. Hearts were excised quickly, washed in PBS and weighed. Hearts were either fixed in 10% buffered formalin or snap-frozen in liquid nitrogen.

To insert the minipumps, mice were anaesthetised in an induction chamber using vaporised 5% isoflurane with 2 l/min oxygen and placed on a heated mat in the prone position. For analgesia, buprenorphine (Vetergesic, Ceva Animal Health Ltd.) (0.05 mg/kg, diluted in sterile PBS) was administered subcutaneously. Anaesthesia was maintained using 2.5% isoflurane in 2 l/min oxygen using a nose cone. The fur covering the mid-scapular region was removed using an electric razor and the area was sterilised with HIBISCRUB^®^ (VioVet). A 2 cm incision was made at the mid-scapular region and blunt dissection generated a pocket towards the lower-left flank of the mouse for the minipump(s) to be inserted. The wound was closed with two simple interrupted sutures using polypropylene 4-0 thread (Prolene, Ethicon) and sterilised with HIBISCRUB^®^.

Echocardiography was performed with a Vevo 2100^TM^ (Fujifilm Visualsonics) using a 38 MHz MS400 transducer. Echocardiograms were collected before minipump surgery and at 3 and 7 d. Anaesthesia used vaporised isoflurane (5% with 1 l/min oxygen for induction; 1.5% in 1 l/min oxygen for maintenance). Mice were placed on a heating physiological monitoring stage in a supine position, and heart rate, respiration rate and body temperature were monitored. Chest fur was removed, pre-warmed ultrasound gel was applied to the chest and the transducer lowered into the gel until a clear image was obtained. Imaging was completed within 10-15 min. Mice were recovered singly and transferred to the home cage.

Speckle tracking (strain) analysis used B-mode images of long axis views of the heart. Analysis was performed across two cardiac cycles starting from the R peak after a breath had been taken so that tracking was not interrupted by movement of the heart associated with the breath. VevoStrain software was used to obtain global measures of cardiac dimensions and function and for segmental analysis. Global strain measurements and endocardial data for each of 6 segments (derived from 9 individual points) around the LV were imported into Excel. For segmental analysis, measurements were obtained for radial and longitudinal peak displacement, velocity, strain and strain rate each representing the average of the two cycles.

### Histological analysis

Hearts were fixed with 10% formalin, immersed in 70% ethanol, embedded in paraffin, sectioned at 10 μm and stained using kits for H&E (Sigma) or Masson’s trichrome (Polysciences). Sections were also stained for picrosirius red (PSR). Sections were submerged in Weigert’s hematoxylin, washed, stained in picrosirius red (1 g/l Sirus red in saturated aqueous picric acid, 60 min), and differentiated in 0.5% acetic acid. Images were captured digitally using a Nikon slide scanner. For analysis of myocyte cross-sectional area, cells stained by H&E within the LV (excluding endocardial regions) with a central, single nucleus in clear cross-section were chosen at random and outline traced using NDP.view2 software (Hamamatsu). To assess interstitial and fibrosis at the percentage level, Massons-trichrome and PSR images of the entire LV were exported and the collagen fraction calculated as the ratio between the sum of the total area of fibrosis (blue or red colour) to the sum of the total tissue area (including the myocyte area) for the entire image using Image-J. For perivascular fibrosis, the degree of fibrosis was scored (0.5, small amount of fibrosis around a single vessel; 1, small amount of fibrosis around 2 or more vessels; 1.5, significant fibrosis around a single vessel; 2, significant fibrosis around 2 or more vessels; 2.5, extensive fibrosis around 1 vessel; 3, extensive fibrosis around 2 or more vessels; 3.5 extensive fibrosis around a single vessel penetrating into the myocardium with extensive fibrosis around other vessels; 4, extensive fibrosis around 2 or more vessels penetrating into the myocardium.

### Western blotting

Heart powders (10-20 mg) were homogenised in 6 vol extraction buffer [20 mM Tris pH 7.5, 1 mM EDTA, 10% (v/v) glycerol, 1% (v/v) Triton X-100, 100 mM KCl, 5 mM NaF, 0.2 mM Na_3_VO_4_, 5 mM MgCl_2_, 0.05% (v/v) 2-mercaptoethanol, 10 mM benzamidine, 0.2 mM leupeptin, 0.01 mM trans-epoxy succinyl-L-leucylamido-(4-guanidino)butane, 0.3 mM phenylmethylsulphonyl fluoride, 4 µM microcystin] and extracted on ice with intermittent vortex mixing (10 min). Samples were centrifuged (10,000 × g, 10 min, 4°C), the supernatants were removed, a sample was taken for protein assay and the rest boiled with 0.33 vol sample buffer (300 mM Tris-HCl pH 6.8, 10% (w/v) SDS, 13% (v/v) glycerol, 130 mM dithiothreitol, 0.2% (w/v) bromophenol blue). Protein concentrations were determined against BSA standards by BioRad Bradford assay using a 1/5 dilution (v/v) in H_2_O of the dye reagent concentrate.

Proteins (40 µg for Pkn1 and Pkn2; 10 µg for Gapdh), were separated by SDS-PAGE (200 V) using 8% (for Pkn1/2), or 12% (Gapdh) polyacrylamide resolving gels with 6% stacking gels . Samples were electrophoresed until the dye front reached the bottom of the gel (∼50 min). Proteins were transferred to nitrocellulose using a BioRad semi-dry transfer cell (10 V, 60 min). Blots were incubated (15 min) with 5% (w/v) non-fat milk powder in Tris-buffered saline (20 mM Tris-HCl pH 7.5, 137 mM NaCl) containing 0.1% (v/v) Tween 20 (TBST) to reduce non-specific binding, then incubated overnight at 4°C with primary antibodies in TBST containing 5% (w/v) BSA. Blots were washed with TBST (3 × 5 min, 21°C), incubated with horseradish peroxidase-conjugated goat anti-rabbit immunoglobulin antibodies (Dako; catalogue no. P0449; 1/5000 dilution) in TBST containing 1% (w/v) non-fat milk powder (60 min, 21°C) and then washed again in TBST (3 × 5 min, 21°C). Rabbit polyclonal antibodies to PKN2 (Cat. no. 2612; 1/750 dilution), phosphorylated PKN1/2 (Cat. no. 2611; 1/750 dilution) and Gapdh (Cat. No. 2118; 1/1000 dilution) were from Cell Signaling Technology. Antibodies to PKN1 were from BD Transduction Laboratories (Cat. no. 610686; 1/750 dilution). Bands were detected by enhanced chemiluminescence using ECL Prime and captured with an ImageQuant LAS4000 system (Cytiva). Densitometry was performed using ImageQuant TL 8.1 software (GE Healthcare).

### RT-qPCR

Samples for all mice were processed simultaneously. Heart powders (10-15 mg) were weighed into safelock Eppendorf tubes and kept on dry ice. RNA Bee (AMS Biotechnology Ltd) was added (1 ml) and samples homogenised on ice using a pestle. RNA was prepared according to the manufacturer’s instructions and dissolved in nuclease-free water. The purity was assessed from the A_260_/A_280_ measured using an Implen NanoPhotometer (values were 1.8–2.0) and concentrations determined from the A_260_. Total RNA was reverse transcribed to cDNA using High Capacity cDNA Reverse Transcription Kits with random primers (Applied Biosystems). qPCR was performed using a StepOnePlus Real-Time PCR system (ThermoFisher Scientific) using 1/40 of the cDNA produced. Optical 96-well reaction plates were used with iTaq Universal SYBR Green Supermix (Bio-Rad Laboratories Inc.) according to the manufacturer’s instructions. See **Supplementary Table S6** for primer sequences. Results were normalized to *Gapdh*, and relative quantification was obtained using the ΔCt (threshold cycle) method; relative expression was calculated as 2^−ΔΔCt^, and normalised to the mean of vehicle controls in wild-type mice.

### Statistical analysis and data presentation

Data were imported into Microsoft Excel. Data analysis used Microsoft Excel with GraphPad Prism 11.02 for statistical analysis and data presentation. A Rout outliers test (Q=1%) was used removing a maximum of 1 of observation per condition. Shapiro-Wilks’ tests were used to confirm normality for all data. Multiple t tests with Holm-Sidak’s multiple comparisons test were used to assess mRNA expression of *Pkn1*, *Pkn2* and *Pkn3* mRNAs in mice treated with vehicle vs AngII at 1, 3 and 7 d. To compare baseline cardiac function/dimensions between wild-type, Pkn2Het and Pkn1/3DKO mice, statistical analysis used unpaired one-way ANOVA with Tukey’s post-test. For effects of AngII, statistical analysis used unpaired two-tailed 2-way ANOVA with Tukey’s multiple comparison post-tests comparing treatments (vehicle, AngII) at 3 and 7 d and genotypes (wild-type, Pkn2Het, Pkn1/3DKO) as appropriate and indicated in the Figure Legends. Images from proprietary software were exported as .tif or .jpg files and cropped for presentation with Adobe Photoshop CC maintaining the original relative proportions.

### Data Availability

The data are available from the authors on request.

### Funding

J.J.T.M. and P.J.P. were supported by the Francis Crick Institute which receives it score funding from Cancer Research UK (CRUK) (FC001130), the UK Medical Research Council (MRC) (FC001130), and the Wellcome Trust (FC001130).

### Selected abbreviations

AngII,: angiotensin II;
CO,: cardiac output;
CSA,: cross-sectional area;
DKO,: double-knockout;
EDV,: end diastolic volume;
ESV,: end systolic volume;
FS,: fractional shortening;
GLS,: global longitudinal strain;
HFpEF,: heart failure with preserved ejection fraction;
HPA,: Human Protein Atlas;
LV,: left ventricle;
LVEDM,: LV end diastolic mass;
PKN,: protein kinase N;
VSMCs,: vascular smooth muscle cells.

