## Supplementary Tables and Figures for "PKNs are required for cardiac hypertrophy induced by angiotensin II in mice"

**Supplementary material**

Susanna TE Cooper[1], Jacqueline JT Marshall[2], Peter J Parker[2], Angela Clerk[3]

[1] Institute of Developmental and Regenerative Medicine, Department of Physiology, Anatomy and Genetics, University of Oxford, UK.
[2] Protein Phosphorylation Laboratory, Francis Crick Institute, 1 Midland road, London NW1 1AT, UK
[3] School of Biological Sciences, University of Reading, Whiteknights Campus, Reading, RG6 2AS, UK

**Supplementary Table S1. Human Protein Atlas data for PKN isoform expression in cardiac cells.**

**Supplementary Table S2. Echocardiography of wild-type, Pkn2Het vs Pkn1/3DKO mouse hearts at baseline: global strain**

**Supplementary Table S3. Echocardiography of wild-type, Pkn2Het vs Pkn1/3DKO mouse hearts at baseline: segmental strain. A**, Radial peak values. **B**, Longitudinal (Long) peak values.

**Supplementary Table S4. Echocardiography of hearts from wild-type, Pkn2Het vs Pkn1/3DKO mice treated with vehicle or 0.8 mg/kg/d AngII: global strain.**

**Supplementary Table S5. Mouse body weights**

**Supplementary S6. qPCR primers.**

**Supplementary spreadsheet SS1. Human Protein Atlas data for PKN isoforms and TMEME119 expression in all tissues.**

**Supplementary Figure S1. HPA mRNA expression data for PKNs in vascular smooth muscle cells and adipocytes in different tissues.**

**Supplementary Figure S2. HPA mRNA expression data for PKNs in different cells in different tissues.**

**Supplementary Table S1. Single cell mRNA data for expression of PKN isoforms in heart muscle (Human Protein Atlas version 25.0).** Normalised data are provided (nCPM).

| Population group | Cell type | PKN1 | PKN2 | PKN3 |
|---|---|---|---|---|
| c-0 | Cardiomyocytes | 12.2 | 141.9 | 1.4 |
| c-1 | Cardiomyocytes | 12.4 | 104.8 | 2.2 |
| c-2 | Cardiomyocytes | 11.8 | 119.9 | 1.4 |
| c-3 | Cardiomyocytes | 13 | 110.7 | 2 |
| c-5 | Cardiomyocytes | 11 | 125.4 | 1.5 |
| c-9 | Cardiomyocytes | 3.5 | 113 | 0 |
| c-10 | Cardiomyocytes | 10.7 | 114.9 | 0 |
| c-12 | Cardiomyocytes | 15.8 | 102.6 | 0 |
| c-4 | Fibroblasts | 13.5 | 99.4 | 2.5 |
| c-11 | Epicardial cells | 15.9 | 63.7 | 2.3 |
| c-14 | Adipocytes | 2.5 | 183.1 | 0 |
| c-7 | Vascular endothelial cells | 28.7 | 92.3 | 44 |
| c-8 | Pericytes | 16.4 | 117.9 | 3 |
| c-17 | Pericytes | 9.9 | 49.4 | 0 |
| c-13 | Vascular smooth muscle cells | 0 | 60.3 | 0 |
| c-6 | Macrophages | 18.8 | 131.4 | 1.9 |
| c-15 | T-cells | 12 | 95.7 | 12 |
| c-16 | Mast cells | 14.7 | 73.4 | 7.3 |

**Supplementary Table S2. Echocardiography of wild-type, Pkn2Het vs Pkn1/3DKO mouse hearts at baseline: global strain.** Analysis used long-axis views of the left ventricle (LV). EDV, end diastolic volume; ESV, end systolic volume; SV, stroke volume; FS, fractional shortening; CO, cardiac output; GLS, global longitudinal strain; EF, ejection fraction; LVEDM, LV end diastolic mass.

| | **Wild-type (n=16)** | | **Pkn2Het (n=15)** | | **Pkn1/3DKO (n=11)** | |
|---|---|---|---|---|---|---|
| | **Mean** | **SD** | **Mean** | **SD** | **Mean** | **SD** |
| **EDV (µl)** | 49.12 | 6.94 | 52.27 | 6.87 | 56.28 | 7.24 |
| **ESV (µl)** | 22.41 | 4.19 | 23.11 | 5.37 | 25.65 | 3.83 |
| **SV (µl)** | 26.71 | 5.32 | 29.16 | 3.83 | 30.63 | 4.85 |
| **FS (%)** | 26.24 | 5.30 | 29.50 | 6.08 | 27.35 | 2.94 |
| **CO (ml/min)** | 12.65 | 2.99 | 14.48 | 2.88 | 13.45 | 2.49 |
| **GLS (%)** | -19.51 | 2.81 | -18.57 | 3.65 | -18.60 | 2.36 |
| **EF (%)** | 53.89 | 6.24 | 55.86 | 6.62 | 54.18 | 4.26 |
| **LVEDM (mg)** | 62.93 | 6.84 | 63.22 | 6.29 | 62.87 | 7.05 |
| **Heart rate (bpm)** | 473 | 46 | 495 | 60 | 438 | 26 |

**Supplementary Table S3. Echocardiography of wild-type, Pkn2Het vs Pkn1/3DKO mouse hearts at baseline: segmental strain.** Analysis used long-axis views of the left ventricle (LV). **A**, Radial peak values. **B**, Longitudinal (Long) peak values.

| **A. Radial peak values** | | **Wild-type (n=16)** | | **Pkn2Het (n=15)** | | **Pkn1/3DKO (n=11)** | | **2-way ANOVA Tukey's post-test** | |
|---|---|---|---|---|---|---|---|---|---|
| | Segment | Mean | SD | Mean | SD | Mean | SD | WT vs 2H | WT vs 1/3 |
| **Radial displacement (mm)** | 1 | 0.55 | 0.14 | 0.65 | 0.12 | 0.62 | 0.13 | **0.037** | 0.284 |
| **Radial displacement (mm)** | 2 | 0.55 | 0.11 | 0.63 | 0.13 | 0.57 | 0.17 | 0.066 | 0.780 |
| **Radial displacement (mm)** | 3 | 0.37 | 0.09 | 0.40 | 0.10 | 0.31 | 0.09 | 0.755 | 0.322 |
| **Radial displacement (mm)** | 6 | 0.19 | 0.08 | 0.20 | 0.11 | 0.23 | 0.07 | 0.944 | 0.595 |
| **Radial displacement (mm)** | 5 | 0.26 | 0.08 | 0.24 | 0.06 | 0.26 | 0.07 | 0.895 | 0.997 |
| **Radial displacement (mm)** | 4 | 0.42 | 0.14 | 0.41 | 0.10 | 0.44 | 0.08 | 0.888 | 0.884 |
| **Radial velocity (cm/s)** | 1 | 1.81 | 0.41 | 2.13 | 0.45 | 1.92 | 0.38 | 0.016 | 0.658 |
| **Radial velocity (cm/s)** | 2 | 1.78 | 0.31 | 2.00 | 0.38 | 1.78 | 0.41 | 0.149 | 0.999 |
| **Radial velocity (cm/s)** | 3 | 1.22 | 0.28 | 1.43 | 0.45 | 1.14 | 0.35 | 0.168 | 0.819 |
| **Radial velocity (cm/s)** | 6 | 0.61 | 0.22 | 0.70 | 0.35 | 0.68 | 0.12 | 0.735 | 0.855 |
| **Radial velocity (cm/s)** | 5 | 0.74 | 0.22 | 0.71 | 0.23 | 0.76 | 0.19 | 0.946 | 0.993 |
| **Radial velocity (cm/s)** | 4 | 1.17 | 0.41 | 1.17 | 0.37 | 1.21 | 0.13 | 0.772 | 0.952 |
| **Radial strain (%)** | 1 | 30.97 | 8.15 | 34.86 | 7.93 | 32.80 | 8.53 | 0.404 | 0.845 |
| **Radial strain (%)** | 2 | 37.01 | 10.47 | 40.08 | 7.86 | 38.50 | 10.70 | 0.569 | 0.895 |
| **Radial strain (%)** | 3 | 26.23 | 10.89 | 25.35 | 6.59 | 20.67 | 9.48 | 0.954 | 0.214 |
| **Radial strain (%)** | 6 | 23.51 | 10.16 | 21.12 | 6.90 | 23.99 | 7.37 | 0.712 | 0.988 |
| **Radial strain (%)** | 5 | 37.18 | 5.96 | 35.06 | 9.32 | 38.51 | 5.57 | 0.765 | 0.915 |
| **Radial strain (%)** | 4 | 38.03 | 9.06 | 35.80 | 8.01 | 38.51 | 6.80 | 0.742 | 0.776 |
| **Radial strain rate (1/s)** | 1 | 8.11 | 2.27 | 9.22 | 2.19 | 8.16 | 1.94 | 0.351 | 0.998 |
| **Radial strain rate (1/s)** | 2 | 9.79 | 2.25 | 10.07 | 1.81 | 9.08 | 1.96 | 0.936 | 0.693 |
| **Radial strain rate (1/s)** | 3 | 7.72 | 2.64 | 8.35 | 1.59 | 7.40 | 2.69 | 0.712 | 0.927 |
| **Radial strain rate (1/s)** | 6 | 6.58 | 2.20 | 6.63 | 2.07 | 7.18 | 1.92 | 0.998 | 0.771 |
| **Radial strain rate (1/s)** | 5 | 9.16 | 1.66 | 8.69 | 2.26 | 9.32 | 1.12 | 0.828 | 0.981 |
| **Radial strain rate (1/s)** | 4 | 9.73 | 2.94 | 9.49 | 2.83 | 9.95 | 2.41 | 0.953 | 0.966 |

| **B. Longitudinal peak values** | | **Wild-type (n=16)** | | **Pkn2Het (n=15)** | | **Pkn1/3DKO (n=11)** | | **2-way ANOVA Tukey's post-test** | |
|---|---|---|---|---|---|---|---|---|---|
| | Segment | Mean | SD | Mean | SD | Mean | SD | WT vs 2H | WT vs 1/3 |
| **Long displacement (mm)** | 1 | 0.49 | 0.11 | 0.47 | 0.12 | 0.47 | 0.15 | 0.924 | 0.827 |
| **Long displacement (mm)** | 2 | 0.21 | 0.08 | 0.16 | 0.04 | 0.21 | 0.06 | 0.275 | 0.988 |
| **Long displacement (mm)** | 3 | 0.06 | 0.05 | 0.05 | 0.02 | 0.07 | 0.06 | 0.977 | 0.890 |
| **Long displacement (mm)** | 6 | 0.03 | 0.01 | 0.02 | 0.02 | 0.03 | 0.01 | 0.961 | 0.998 |
| **Long displacement (mm)** | 5 | 0.19 | 0.06 | 0.16 | 0.07 | 0.17 | 0.06 | 0.564 | 0.775 |
| **Long displacement (mm)** | 4 | 0.46 | 0.13 | 0.45 | 0.16 | 0.47 | 0.12 | 0.851 | 0.997 |
| **Long velocity (cm/s)** | 1 | 1.56 | 0.50 | 1.38 | 0.45 | 1.37 | 0.48 | 0.308 | 0.342 |
| **Long velocity (cm/s)** | 2 | 0.72 | 0.23 | 0.62 | 0.19 | 0.62 | 0.15 | 0.801 | 0.720 |
| **Long velocity (cm/s)** | 3 | 0.31 | 0.19 | 0.26 | 0.12 | 0.31 | 0.16 | 0.917 | >0.999 |
| **Long velocity (cm/s)** | 6 | 0.25 | 0.09 | 0.22 | 0.12 | 0.23 | 0.08 | 0.975 | 0.991 |
| **Long velocity (cm/s)** | 5 | 0.84 | 0.32 | 0.59 | 0.20 | 0.63 | 0.22 | 0.101 | 0.249 |
| **Long velocity (cm/s)** | 4 | 1.72 | 0.69 | 1.42 | 0.53 | 1.49 | 0.40 | 0.046 | 0.195 |
| **Long strain (%)** | 1 | -22.82 | 6.14 | -20.00 | 5.56 | -22.93 | 3.93 | 0.255 | 0.998 |
| **Long strain (%)** | 2 | -20.47 | 3.53 | -19.15 | 5.51 | -19.19 | 6.65 | 0.741 | 0.787 |
| **Long strain (%)** | 3 | -20.41 | 5.26 | -22.07 | 8.01 | -17.41 | 3.54 | 0.989 | 0.273 |
| **Long strain (%)** | 6 | -14.24 | 6.08 | -14.80 | 6.72 | -14.83 | 2.32 | 0.947 | 0.950 |
| **Long strain (%)** | 5 | -16.30 | 3.66 | -15.32 | 3.79 | -15.90 | 2.12 | 0.849 | 0.977 |
| **Long strain (%)** | 4 | -19.60 | 4.98 | -16.67 | 5.01 | -21.41 | 5.82 | **0.043** | 0.990 |
| **Long strain rate (1/s)** | 1 | -8.44 | 3.01 | -7.33 | 2.96 | -8.30 | 1.86 | 0.393 | 0.987 |
| **Long strain rate (1/s)** | 2 | -7.05 | 2.16 | -6.52 | 2.42 | -6.35 | 3.28 | 0.804 | 0.733 |
| **Long strain rate (1/s)** | 3 | -6.89 | 1.96 | -8.55 | 4.50 | -6.34 | 1.87 | 0.691 | 0.826 |
| **Long strain rate (1/s)** | 6 | -5.09 | 2.25 | -5.36 | 2.79 | -4.82 | 1.22 | 0.946 | 0.956 |
| **Long strain rate (1/s)** | 5 | -5.52 | 2.07 | -4.67 | 1.39 | -5.10 | 1.08 | 0.576 | 0.891 |
| **Long strain rate (1/s)** | 4 | -7.35 | 2.98 | -6.09 | 2.17 | -7.03 | 2.72 | 0.304 | 0.934 |

**Supplementary Table S4. Echocardiography of hearts from wild-type, Pkn2Het vs Pkn1/3DKO mice treated with vehicle or 0.8 mg/kg/d AngII: global strain.** Analysis used long-axis views of the left ventricle (LV). **A**, Echocardiograms collected at 3 d. **B**, Echocardiograms collected at 7 d.

| A. 3 d | Wild-type | | | | Pkn2Het | | | | Pkn1/3DKO | | | |
|---|---|---|---|---|---|---|---|---|---|---|---|---|
| | Vehicle (n=8) | | AngII (n=6) | | Vehicle (n=8) | | AngII (n=7) | | Vehicle (n=5) | | AngII (n=6) | |
| | Mean | SD | Mean | SD | Mean | SD | Mean | SD | Mean | SD | Mean | SD |
| **EDV (µl)** | 48.00 | 6.75 | 42.43 | 9.91 | 54.78 | 10.44 | 45.39 | 11.39 | 55.94 | 4.12 | 38.85 | 11.54 |
| **ESV (µl)** | 19.87 | 4.42 | 17.58 | 6.48 | 24.71 | 6.02 | 18.34 | 5.30 | 23.29 | 4.20 | 13.65 | 5.21 |
| **SV (µl)** | 28.13 | 3.23 | 24.84 | 5.21 | 30.08 | 4.85 | 27.05 | 7.22 | 32.66 | 1.86 | 25.20 | 7.13 |
| **FS (%)** | 31.52 | 5.12 | 31.87 | 8.85 | 26.12 | 4.51 | 29.31 | 1.78 | 32.48 | 4.59 | 36.41 | 5.05 |
| **CO (ml/min)** | 13.59 | 1.91 | 13.21 | 2.74 | 14.79 | 1.88 | 13.65 | 4.40 | 15.49 | 2.07 | 12.54 | 3.77 |
| **GLS (%)** | - 20.61 | 2.27 | - 20.51 | 8.14 | - 18.91 | 1.33 | - 19.21 | 5.57 | -19.93 | 0.90 | - 22.85 | 2.36 |
| **EF (%)** | 58.68 | 4.93 | 59.37 | 10.53 | 54.93 | 3.80 | 59.04 | 6.78 | 58.32 | 4.91 | 64.91 | 5.17 |
| **LVEDM (mg)** | 61.65 | 5.53 | 77.62 | 6.06 | 62.31 | 8.30 | 71.46 | 5.42 | 65.78 | 5.22 | 80.86 | 8.51 |
| **Heart rate (bpm)** | 483 | 30 | 540 | 104 | 494 | 24 | 497 | 61 | 474 | 57 | 497 | 29 |

| B. 7 d | Wild-type | | | | Pkn2Het | | | | Pkn1/3DKO | | | |
|---|---|---|---|---|---|---|---|---|---|---|---|---|
| | Vehicle (n=8) | | AngII (n=6) | | Vehicle (n=8) | | AngII (n=7) | | Vehicle (n=5) | | AngII (n=6) | |
| | Mean | SD | Mean | SD | Mean | SD | Mean | SD | Mean | SD | Mean | SD |
| **EDV (µl)** | 54.97 | 5.88 | 42.30 | 10.49 | 51.97 | 4.64 | 44.76 | 9.61 | 57.16 | 8.21 | 44.41 | 8.81 |
| **ESV (µl)** | 24.95 | 4.10 | 16.07 | 5.80 | 23.39 | 4.66 | 19.81 | 2.96 | 24.54 | 4.51 | 18.77 | 6.62 |
| **SV (µl)** | 30.02 | 2.66 | 26.23 | 6.84 | 28.57 | 1.85 | 24.95 | 6.97 | 32.62 | 4.15 | 25.64 | 2.42 |
| **FS (%)** | 26.24 | 3.12 | 34.39 | 7.32 | 29.06 | 4.85 | 26.98 | 5.19 | 30.70 | 3.74 | 31.87 | 7.13 |
| **CO (ml/min)** | 14.15 | 1.16 | 13.73 | 3.91 | 14.50 | 1.39 | 12.82 | 3.19 | 16.28 | 1.81 | 12.27 | 0.66 |
| **GLS (%)** | - 20.34 | 0.97 | - 20.33 | 3.12 | - 19.49 | 1.89 | - 17.71 | 5.09 | -18.76 | 0.58 | - 19.03 | 2.83 |
| **EF (%)** | 54.62 | 3.59 | 61.81 | 7.98 | 55.04 | 5.18 | 54.82 | 4.68 | 56.81 | 2.88 | 58.88 | 8.76 |
| **LVEDM (mg)** | 63.10 | 4.96 | 89.78 | 11.91 | 64.53 | 3.27 | 71.81 | 7.48 | 72.18 | 3.45 | 72.08 | 11.77 |
| **Heart rate (bpm)** | 472 | 26 | 520 | 44 | 508 | 44 | 519 | 43 | 500 | 21 | 482 | 46 |

**Supplementary Table S5. Mouse body weights.** Data are for male mice.

A. Baseline body weights for wild-type and Pkn1/3DKO mice. Statistical analysis used 1-way ANOVA with Tukey's post-test.

| | Mean | SD | n | p (vs wild-type) |
|---|---|---|---|---|
| Wild-type | 28.04 | 1.292 | 16 | |
| Pkn2Het | 28.17 | 2.061 | 15 | 0.838 |
| Pkn1/3DKO | 29.33 | 1.852 | 11 | 0.129 |

B. Body weights for wild-type mice treated with vehicle or 0.8 mg/kg/d AngII. Statistical analysis used 2-way ANOVA with Tukey's post-test.

| | Vehicle | | | AngII | | | p (Veh vs AngII) |
|---|---|---|---|---|---|---|---|
| | Mean | SD | n | Mean | SD | n | |
| Baseline | 28.23 | 1.17 | 9 | 27.79 | 1.49 | 7 | 0.584 |
| Post-surgery | 28.47 | 2.07 | 9 | 28.93 | 1.84 | 7 | 0.572 |
| 3 d | 29.24 | 1.32 | 9 | 28.41 | 1.69 | 7 | 0.311 |
| 7 d | 29.28 | 1.60 | 9 | 28.54 | 1.59 | 7 | 0.370 |

C. Body weights for Pkn2Het mice treated with vehicle or 0.8 mg/kg/d AngII. Statistical analysis used 2-way ANOVA with Tukey's post-test.

| | Vehicle | | | AngII | | | P (Veh vs AngII) |
|---|---|---|---|---|---|---|---|
| | Mean | SD | n | Mean | SD | n | |
| Baseline | 28.34 | 1.88 | 8 | 27.97 | 2.39 | 7 | 0.733 |
| Post-surgery | 29.29 | 2.15 | 8 | 28.47 | 2.08 | 7 | 0.447 |
| 3 d | 29.45 | 2.15 | 8 | 28.47 | 1.65 | 7 | 0.363 |
| 7 d | 29.93 | 2.16 | 8 | 28.97 | 1.91 | 7 | 0.375 |

D. Body weights for Pkn1/3DKO mice treated with vehicle or 0.8 mg/kg/d AngII. Statistical analysis used 2-way ANOVA with Tukey's post-test.

| | Vehicle | | | AngII | | | P (Veh vs AngII) |
|---|---|---|---|---|---|---|---|
| | Mean | SD | n | Mean | SD | n | |
| Baseline | 30.06 | 1.16 | 5 | 28.72 | 2.19 | 6 | 0.189 |
| Post-surgery | 30.28 | 1.52 | 5 | 29.45 | 1.74 | 6 | 0.413 |
| 3 d | 30.20 | 1.28 | 5 | 29.28 | 1.62 | 6 | 0.367 |
| 7 d | 30.74 | 1.75 | 5 | 29.63 | 1.62 | 6 | 0.277 |

### Supplementary S6. qPCR primers.

| Gene Symbol | Accession No. | Sense Primer (5’→3’) | Antisense Primer (5’→3’) |
|---|---|---|---|
| Areg | ENSMUSG00000029378 | CGCTTATGGTGGAAACCTCTC | GGTCTTAGGCTCAGGCCATTA |
| Cd4 | ENSMUSG00000023274 | GGTGATGAAAGTGGCTCAG | AGGGTCAGTCTCATCTTGG |
| Cd8a | ENSMUSG00000053977 | CTACTATTTCTGCTCAGTCATCAG | GTAGAGTTCACTTTCTGAAGGAC |
| Col1a1 | ENSMUSG00000001506 | TCGTGGCTTCTCTGGTCTC | CCGTTGAGTCCGTCTTTGC |
| Col3a1 | ENSMUSG00000026043 | GGAACCTGGTTTCTTCTCACC | TAGGACTGACCAAGGTGGCT |
| Cspg4 (Ng2) | ENSMUSG00000032911 | TTGGCTACGTGAAGATAGGG | AGCACGATGACTCTGAGACC |
| Ddr2 | ENSMUSG00000026674 | GCACTTGGTGAATTAATTAGAATC CTG | GGACAACTAAATGGTCCCTCCC |
| Ereg | ENSMUSG00000029377 | CACCGAGAAAGAAGGATGGAG | TGTGCTGATAACTGCCTGTAG |
| Fn1 | ENSMUSG00000026193 | AAGAGGACGTTGCAGAGCTA | AGACACTGGAGACACTGACTAA |
| Gapdh | ENSMUSG00000057666 | TCACCACCATGGAGAAGGC | GCTAAGCAGTTGGTGGTGCA |
| Hbegf | ENSMUSG00000024486 | CGGGGAGTGCAGATACCTG | TTCTCCACTGGTAGAGTCAGC |
| Ifna | ENSMUSG00000095498 | TGTGACCTTCCTCAGACTC | TCATTTGTACCAGGAGTGTC |
| Ifng | ENSMUSG00000055170 | CACCTGATTACTACCTTCTTCAG | GTTGTTGACCTCAAACTTGG |
| Il6 | ENSMUSG00000025746 | TCCATCCAGTTGCCTTCTTG | GGTCTGTTGGGAGTGGTATC |
| Itgam (Cd11b) | ENSMUSG00000030786 | CAGATCAACAATGTGACCGTATGG G | CATCATGTCCTTGTACTGCCGCTTG |
| Itgax (Cd11c) | ENSMUSG00000030789 | CTGGATAGCCTTTCTTCTGCTG | GCACACTGTGTCCGAACTC |
| Myh7 | ENSMUSG00000053093 | CATGCCAACCGTATGGCTG | GTTCCACGATGGCGATGTTC |
| Nppa | ENSMUSG00000041616 | GATGGATTTCAAGAACCTGCTAGA | CTTCCTCAGTCTGCTCACTCA |
| Nppb | ENSMUSG00000029019 | TCCAGCAGAGACCTCAAAATTC | CAGTGCGTTACAGCCCAAA |
| Nrg1 | ENSMUSG00000062991 | GAGAACACCCAAGTCAGGA | GATGTAGATGTGGAAAGTTTAGGAG |
| Pecam1 (Cd31) | ENSMUSG00000020717 | CTGCCAGTCCGAAAATGGAAC | CTTCATCCACCGGGGCTATC |
| Pkn1 | ENSMUSG00000057672 | ATCCGGCACACTGGAGACA | TGGTGGATTCATGGGTTGGAG |
| Pkn2 | ENSMUSG00000004591 | CTGTGCCTGCAACAGTGC | CTGTAGAGTCACTGGCCGGA |
| Pkn3 | ENSMUSG00000026785 | CTTGGGAACCCCCAACCTTT | CAGATCTTTGCAGCCCAGGA |
| Postn | ENSMUSG00000027750 | TTCCTCTCCTGCCCTTATATGC | CCTGATCCCGACCCCTGAT |
| Ptprc (Cd45) | ENSMUSG00000026395 | GGGTTGTTCTGTGCCTTGTT | CTGGACGGACACAGTTAGCA |
| Sele | ENSMUSG00000026582 | ACTGAAGGGATCAAGAAGAC | CATTTCACAGCTGAACACG |
| Tagln | ENSMUSG00000032085 | GACTGCACTTCTCGGCTCAT | CCGAAGCTACTCTCCTTCCA |

**Supplementary Figure S1. HPA mRNA expression data for PKNs in vascular smooth muscle cells (A) and adipocytes (B) in different tissues.** Identifying cell numbers are provided in parentheses.

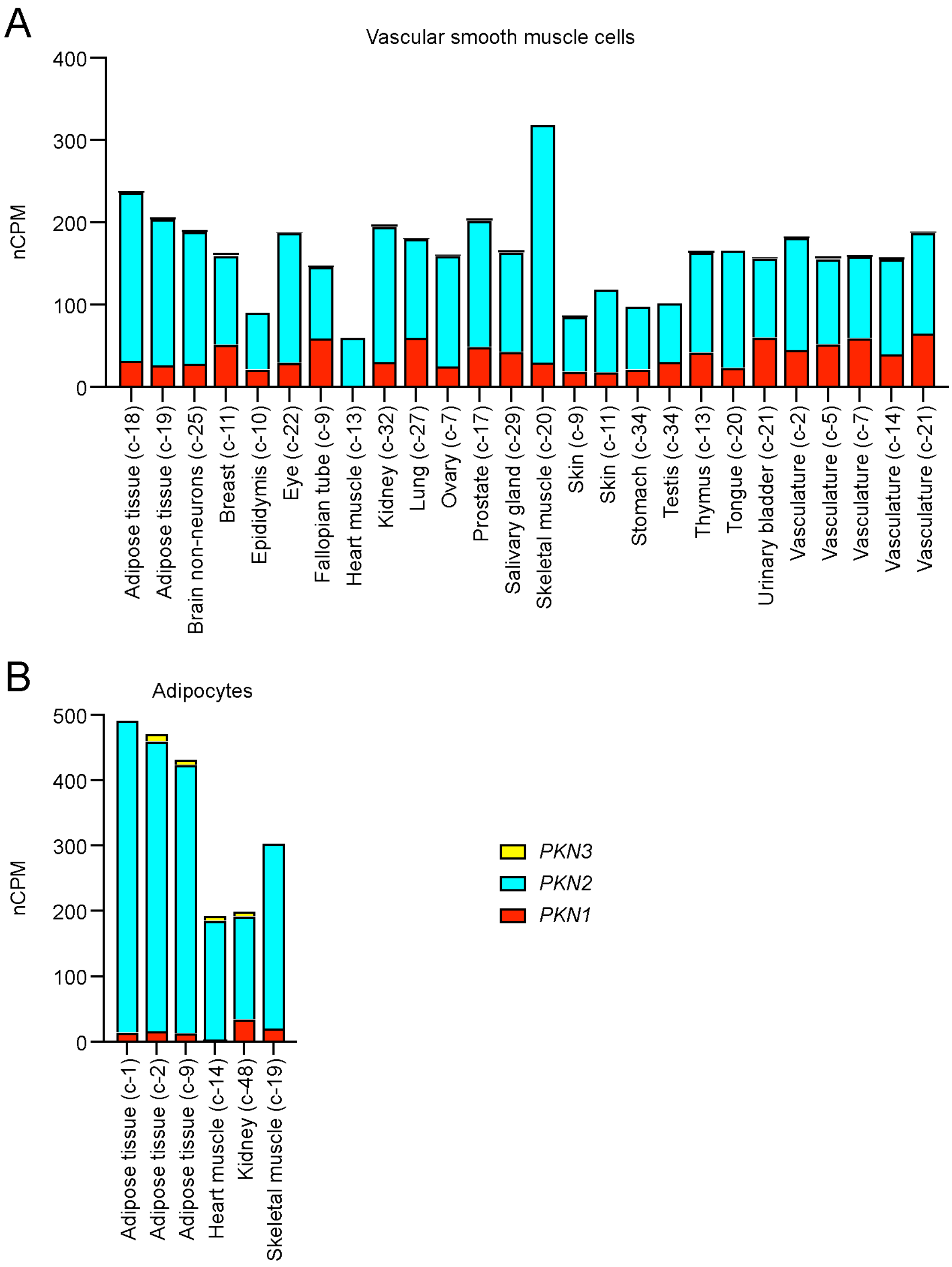

**Supplementary Figure S2. HPA mRNA expression data for PKNs in different cells in different tissues.** Identifying cell numbers are provided in Supplementary Spreadsheet SS1.

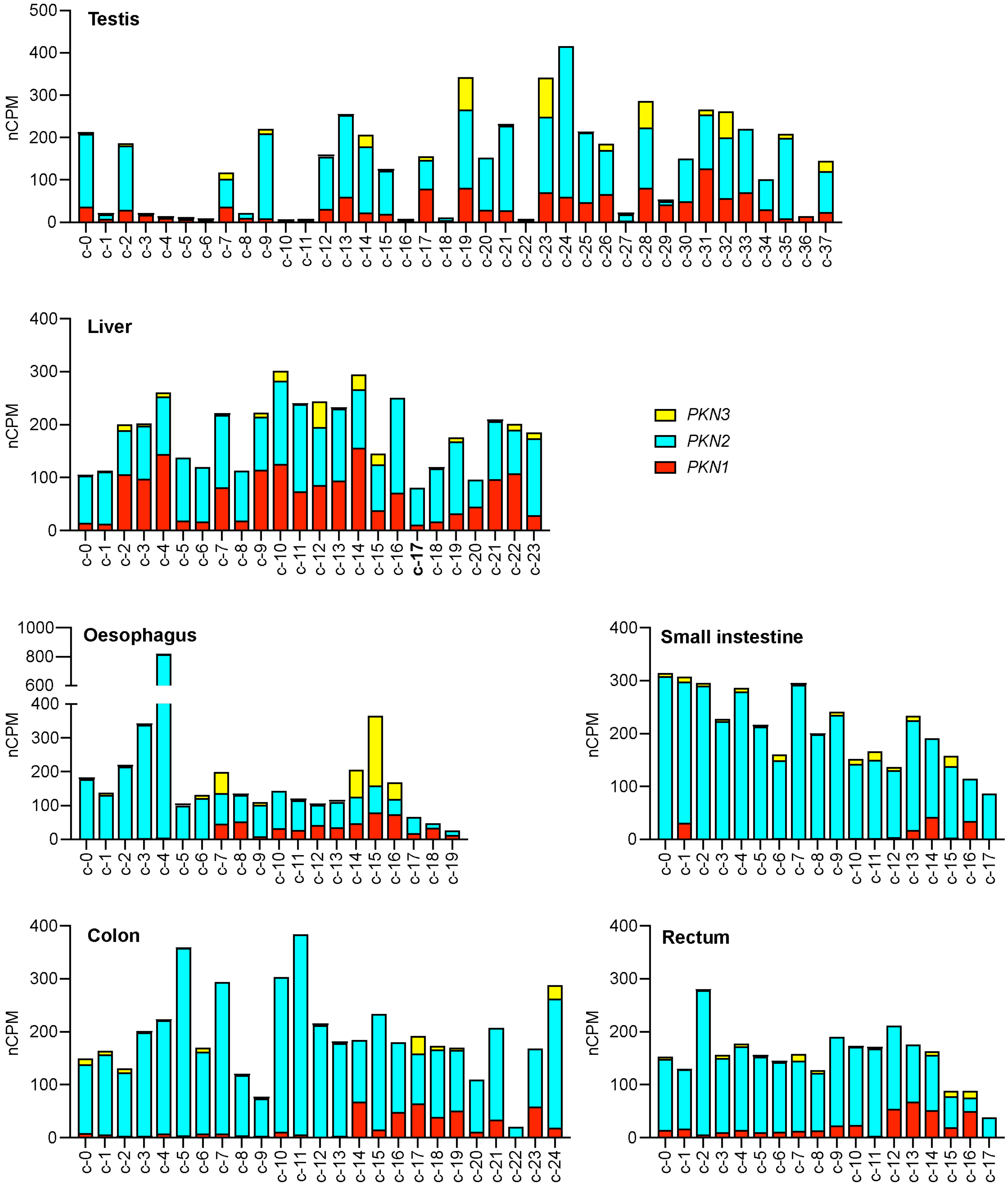
